# ATR kinase inhibitors induce mitochondrial fission in CD8^+^ T cells and impair immune memory *in vivo*

**DOI:** 10.64898/2026.05.25.727628

**Authors:** Frank P. Vendetti, Carina R. Sclafani, Yunqi Zhang, Pinakin Pandya, Yvonne M. Mowery, Su Hyeong Kim, Rachel L. Cumberland, Satishkumar R. Ganakammal, Thomas P. Conrads, Michael J. Calderon, Simon C. Watkins, Bennett Van Houten, Ava Clump, Robert L. Ferris, Jan H. Beumer, Greg M. Delgoffe, Lawrence P. Kane, Christopher J. Bakkenist

## Abstract

The DNA damage response kinase ATR restrains CDK1 activity during S and G2 phases of the cell cycle, confining CDK1-driven processes to mitosis. ATR kinase inhibitors were originally developed to potentiate chemotherapy-induced DNA damage at stalled replication forks and to disrupt DNA damage-induced cell cycle checkpoints. Recent evidence, however, reveals that these inhibitors also disrupt cell cycle organization in cells that have not sustained any DNA damage. We show that ATR kinase inhibitors potently trigger unscheduled mitochondrial fission, causing loss of mitochondrial mass in actively dividing CD8^+^ T cells that persists in memory CD8^+^ T cells. Moreover, ATR inhibition during the peak of CD8^+^ T cell expansion in a mouse model of LCMV Armstrong infection impairs the formation of immune memory. These findings carry significant clinical implications. ATR kinase inhibitors are currently being evaluated in clinical trials in combination with chemotherapy, radiation, and immune checkpoint inhibitors in patients where anti-tumor immune responses are recognized as a determinant of durable response. Our results identify an unexpected consequence of ATR inhibition that disrupts cellular metabolism with broad implications for both preclinical research and clinical application.

## INTRODUCTION

Ataxia telangiectasia and Rad3-related (ATR) is an essential kinase that initiates DNA damage signaling at stalled and collapsed replication forks and single-stranded DNA, ultimately inhibiting cyclin-dependent kinase 1 and 2 (CDK1 and CDK2) and thereby arresting the cell cycle and limiting origin firing (1, 2). ATR kinase inhibitors induce CDK1-dependent origin firing and reduce dNTP biosynthesis in otherwise unperturbed cells, revealing a basal level of homeostatic ATR signaling even in the absence of DNA damaging agents (3-5). Current data support a model in which ATR activity suppresses CDK1 throughout S and G2 phase, while ATR kinase inhibitors relieve this inhibitory signaling, thereby activating CDK1 and triggering premature cell cycle events in S and G2 phase (4, 6). These effects of ATR kinase inhibitors are particularly pronounced in rapidly proliferating CD8^+^ T cells, which complete a full cell cycle in only 4-6 hours (7, 8).

ATR kinase inhibitors were developed to potentiate genotoxic chemotherapies and ceralasertib (AZD6738), berzosertib (VX-970, M6620), elimusertib (BAY 1895344), and tuvusertib (M1774), TCC1727 (NCT07371663), IMP9064 (NCT05269316), and ART0380 (NCT04657068) have advanced to Phase II clinical trials (9-12). In these trials, ATR kinase inhibitors are being evaluated in combination with genotoxic chemotherapy, poly(ADP-ribose) polymerase (PARP) inhibitors, radiotherapy (XRT), and immunotherapy. Substantial preclinical evidence demonstrates that ceralasertib, the most studied ATR kinase inhibitor, sensitizes tumor cells to a range of DNA-damaging agents (13-23).

Preclinical studies also demonstrate that ceralasertib can potentiate anti-tumor immune responses (14, 23-29). ATR inhibition reshapes the immune landscape of both the tumor microenvironment and the periphery, increasing tumor MHC-I expression, reducing tumor PD-L1 expression, and potentiating IFN-I signaling, collectively driving CD8^+^ T cell-dependent anti-tumor responses in mouse models treated with radiotherapy (XRT) (14, 23-25, 30-33). However, ATR inhibition also causes toxicity in proliferating T cells in preclinical models, and causes anemia, myelosuppression, and depletion of circulating proliferating leukocytes in clinical settings (7, 8, 25, 27, 34, 35).

Several studies using syngeneic mouse models of cancer suggest that ceralasertib may potentiate immunologic memory. Two studies found that tumor-bearing mice with a complete response to either XRT plus ceralasertib, or XRT plus ceralasertib plus anti-PD-L1, were resistant to tumor rechallenge in the contralateral flank weeks to months later (23, 25). However, these studies are inconclusive, as neither included control groups in which primary tumors were surgically excised from non-responding animals (vehicle, XRT, or XRT plus anti-PD-L1) prior to rechallenge. In a third study, mice were rechallenged in the contralateral flank on day 20 relative to the start of treatment; secondary tumor growth was significantly slower in mice treated with XRT, ceralasertib, and anti-PD-L1 compared to mice receiving XRT alone or XRT combined with either ceralasertib or anti-PD-L1 individually (24). While this study demonstrated a ceralasertib-specific contribution to secondary tumor control, the rechallenge occurred in mice still bearing the primary tumor and it therefore cannot separate a heightened effector response from a true memory response. To date, no studies have examined the impact of ATR kinase inhibitors on immune memory in a controlled system where the primary response has resolved, the effector population has contracted, and only memory cells persist.

Lymphocytic choriomeningitis virus (LCMV) Armstrong generates a prototypical CD8^+^ T cell response that acutely eliminates viral infection with well-defined kinetics and, importantly, does not induce a robust DNA damage response (36). Following infection with LCMV Armstrong, days 4–8 of clonal expansion are critical for determining terminal effector versus memory precursor cell fates, while the subsequent contraction phase (days 9–30) dictates how many memory cells ultimately survive (37, 38). A previous study with LCMV clone 13, which establishes a chronic viral infection that is not cleared by the CD8^+^ T cell response, showed that treatment with ceralasertib and anti-PD-L1 for seven days starting day eleven days post-infection increased the number of antigen-specific IFN-γ^+^ and IFN-γ^+^ TNF-α^+^ CD8^+^ T cells (27). Ceralasertib also upregulated PD-1 expression on antigen-specific T cells in LCMV clone 13-infected mice, as it does in tumor-bearing mice, rendering them more sensitive to PD-L1 blockade (23, 27). This study did not examine the effects of ATRi on immune memory.

Here we study the impact of ceralasertib (subsequently referred to as ATRi) on CD8^+^ T cells activated *ex vivo* and *in vivo* in response to LCMV Armstrong infection. Remarkably, we show that ceralasertib induces CDK1-dependent phosphorylation of DRP1 in S and G2/M phase cells, triggering mitochondrial fission, reducing mitochondrial mass, and permanently impairing virus-specific CD8^+^ T cell memory pools. These data underscore ATR signaling as a core regulator of the unperturbed cell cycle that temporally confines CDK1 activity to mitosis, and they uncover an exquisitely sensitive temporal requirement for mitochondrial fission during effector expansion in the establishment of CD8^+^ T cell memory.

## RESULTS

### ATRi induces CDK1-dependent DRP1 phosphorylation, mitochondrial fission, and reduced mitochondrial mass and function in *ex vivo* activated CD8^+^ T cells

ATR activity inhibits CDK1, while ATR kinase inhibitors alleviate inhibitory signaling and activate CDK1, causing mitotic events in S and G2 phase (4, 6). DRP1, a cytosolic GTPase and the primary executor of mitochondrial fission, is activated during mitosis by CDK1-mediated phosphorylation at Ser-616 (39, 40). We tested the hypothesis that ATRi would induce CDK1-dependent mitochondrial fission in CD8^+^ T cells as the effects of ATR kinase inhibitors are particularly pronounced in these rapidly proliferating cells (7, 41).

First, we isolated CD8^+^ T cells from wildtype C57BL/6 mice, activated the cells for 24 h *ex vivo* using anti-CD3 and anti-CD28 antibodies and expanded the activated cells for 48 h in IL-2. We then treated the clonally expanding CD8^+^ T cells with 5 µM ATRi for 30 min and assessed phosphorylated DRP1 (pDRP1 S616), along with total DRP1, by immunoblotting. ATRi increased pDRP1 S616 but not total DRP1 protein (Fig 1A). We also examined DRP1 phosphorylation via flow cytometry, using the same pDRP1 S616 primary antibody, Alexa Fluor 647-labeled secondary antibody, and FxCycle Violet to determine DNA content for cell cycle distribution (Fig 1B). ATRi more than doubled the pDRP1 S616 median fluorescence intensity (MFI, normalized to vehicle control) in S and G2/M phase cells (Fig 1C-D).

**Figure 1.**
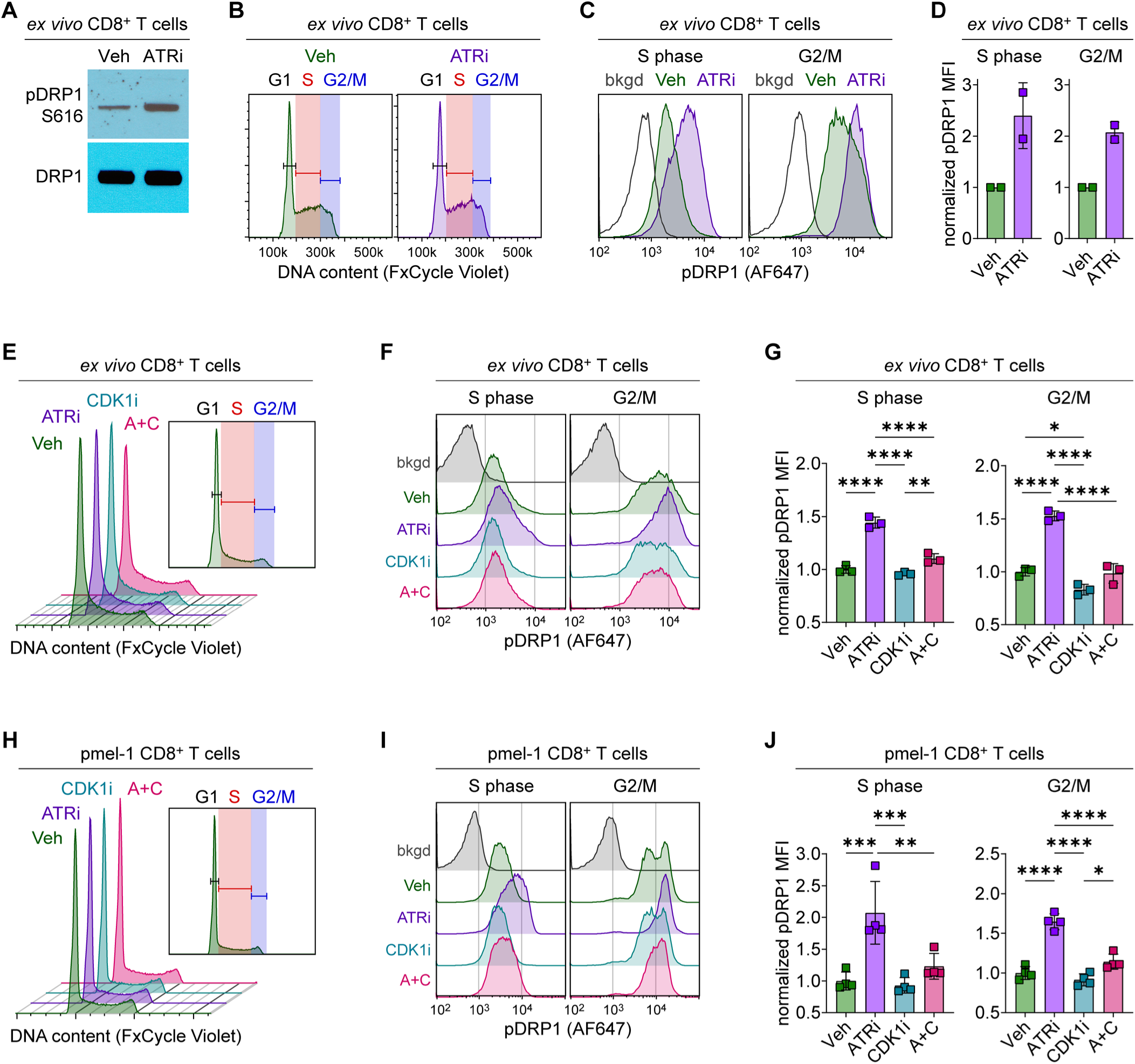
ATRi induces CDK1-dependent DRP1 phosphorylation during S and G2/M phase in clonally expanding CD8^+^ T cells activated *ex vivo*. A-G. CD8^+^ T cells were isolated (via negative selection) from wildtype C57BL/6 spleens and were activated *ex vivo* with plate-bound anti-CD3 (10 µg/mL), soluble anti-CD28 (2 µg/mL), and 50 U/mL IL-2 for 24 h. Activated CD8^+^ T cells were expanded in 50 U/mL IL-2 for 48 h. **A-D**. At 72 h post-activation, CD8^+^ T cells were treated for 30 min with Vehicle (Veh) or 5 µM ATRi and were then immediately lysed or fixed for downstream analyses. **A.** Immunoblots of phosphorylated DRP1 (pDRP1 S616) and total DRP1 in whole cell extracts from lysed CD8^+^ T cells. **B-D.** Flow cytometry analyses of pDRP1 fluorescence (Alex Fluor 647 secondary) and DNA content (FxCycle Violet) in fixed CD8^+^ T cells. **B.** Representative DNA content histograms with cell cycle phase gating indicated. **C.** Representative histograms of pDRP1 fluorescence intensity, and background (bkgd) fluorescence (secondary antibody only control), in S phase and G2/M phase cells. **D.** Quantitation of background-corrected pDRP1 median fluorescence intensity (MFI), normalized to the Veh control, for CD8^+^ T cells in S or G2/M phase. Data combined from two independent experiments, each with one biological replicate (from two pooled spleens). Mean ± SD bars shown. **E-G.** At 72 h post-activation, CD8^+^ T cells were treated for 30 min with Veh or 5 µM ATRi ± 5 µM CDK1i and were then immediately fixed for flow cytometry analyses of pDRP1 fluorescence and DNA content. **E.** Representative DNA content histograms with cell cycle phase gating indicated. **F.** Representative histograms of pDRP1 fluorescence intensity in S phase and G2/M phase cells, with bkgd histograms included. **G.** Quantitation of background-corrected pDRP1 MFI, normalized to the mean of Veh controls, for CD8^+^ T cells in S or G2/M phase. Data from one experiment with three independently isolated and activated CD8^+^ T cell samples. **H-I.** Splenocytes from pmel-1 TCR transgenic mice were activated *ex vivo* with 1 µM gp100 peptide and 50 U/mL IL-2 for 24 h, expanded in 50 U/mL IL-2 for 48 h, treated with Veh or 5 µM ATRi ± 5 µM CDK1i for 30 min, and then stained for flow cytometry analyses of DRP1 phosphorylation and DNA content in pmel-1 CD8^+^ T cells (identified by TCR-β^+^CD8^+^ staining). **H.** Representative DNA content histograms with cell cycle phase gating indicated. **I.** Representative histograms of pDRP1 fluorescence intensity in S phase and G2/M phase cells, with bkgd histograms included. **J.** Quantitation of background-corrected pDRP1 MFI, normalized to the mean of Veh controls, for pmel-1 CD8^+^ T cells in S and G2/M phase. Data from one experiment with four independently harvested and activated pmel-1 spleens**. G, J.** Individual data points with mean ± SD bars shown. *p<0.05, **p<0.01, ***p<0.001 ****p<0.0001 by one-way ANOVA with Tukey’s multiple comparisons test. All significant comparisons shown.

Next, we examined whether the increase in pDRP1 S616 MFI in S and G2/M phase CD8^+^ T cells following ATRi treatment was CDK1-dependent in cells that were activated *ex vivo* and expanded as described above (Fig 1E). ATRi significantly increased pDRP1 S616 MFI in S and G2/M phase CD8^+^ T cells, and these ATRi-induced increases in pDRP1 S616 MFI were almost entirely blocked by addition of the CDK1 inhibitor Ro-3306 (subsequently referred to as CDK1i) (Fig 1F-G). To better mimic physiologic CD8^+^ T cell activation, we examined pDRP1 S616 MFI in antigen peptide-stimulated and clonally expanded CD8^+^ T cells from pmel-1 TCR-transgenic mice. We stimulated splenocytes from pmel-1 mice with gp100 peptide and IL-2 for 24 h and then expanded the splenocytes for 48 h in IL-2. We treated the splenocytes with ATRi and/or CDK1i for 30 min and examined pDRP1 S616 MFI in S and G2/M phase CD8^+^ T cells (identified by TCR-β^+^ CD8^+^ staining and DNA content) (Fig 1H-I). ATRi again significantly increased pDRP1 S616 MFI in S and G2/M phase CD8^+^ T cells, and these ATRi-induced increases were almost entirely blocked by addition of CDK1i (Fig 1I-J).

We also found that the impact of ATRi on pDRP1 S616 is not restricted to CD8^+^ T cells. In HEK 293T treated with 5 µM ATRi and/or 5 µM CDK1i for 30 min, ATRi significantly increased normalized pDRP1 S616 MFI in non-G1 phase (S/G2/M phase) cells, and this increase was blocked by addition of CDK1i (Supplemental Fig S1A). To validate that ATRi impacts mitochondrial fission, we examined mitochondrial networks by microscopy in cells stained with MitoView Fix 640. We chose two adherent cell lines, mouse embryonic fibroblasts (MEF) and HeLa cells (human cervical cancer), that are well-characterized for microscopy. We treated cells for 6 h with 5 µM ATRi, washed out the inhibitor and allowed cells to recover for an additional 18 h, and stained cells with MitoView Fix 640 for the final 2 h of the recovery period. ATRi treatment resulted in more diffuse mitochondrial staining, indicative of smaller and/or more fragmented mitochondria, in both MEF and HeLa cells (Fig 2A-B).

**Figure 2.**
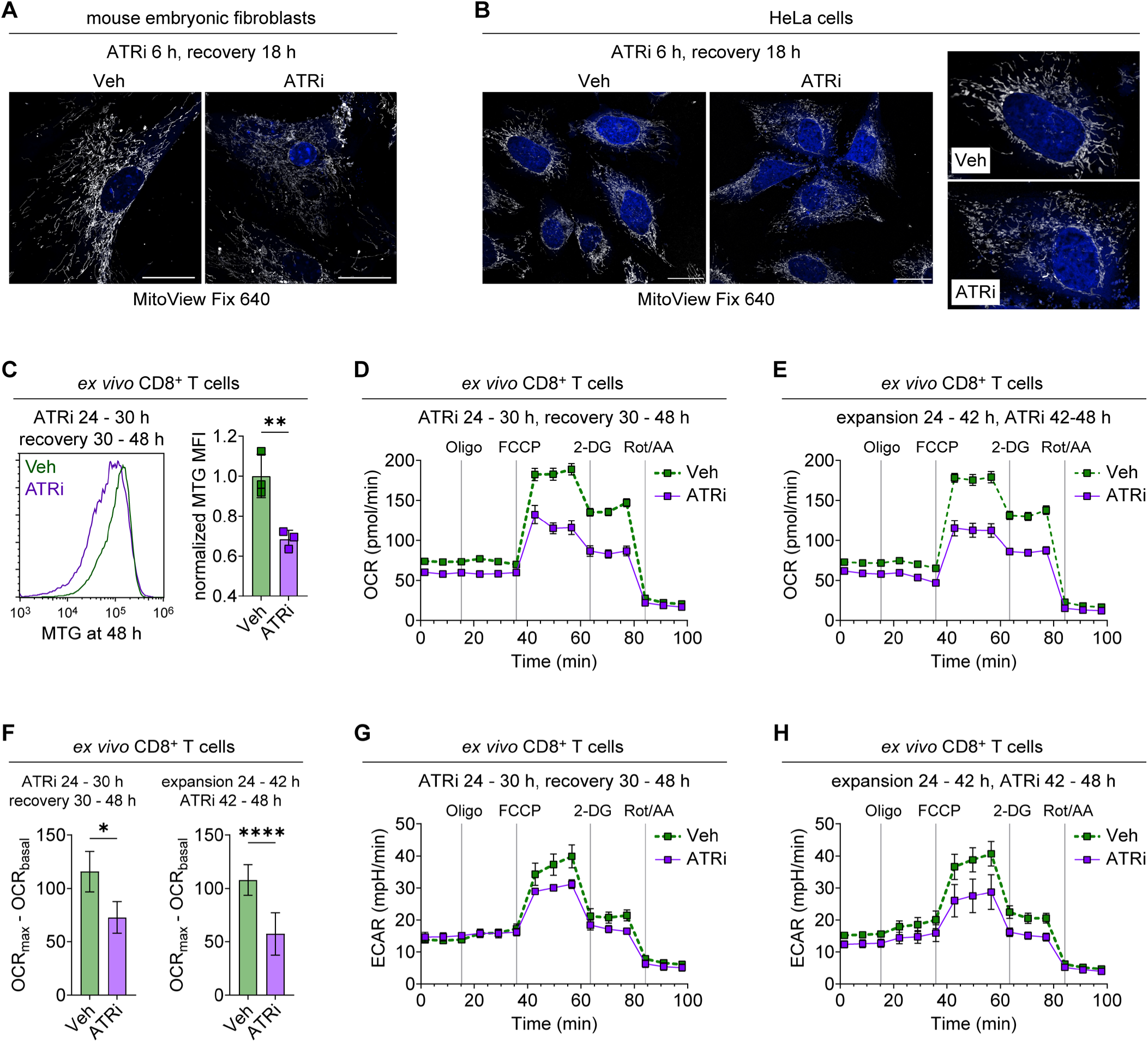
ATRi-induced mitochondrial fission reduces mitochondrial mass and impairs bioenergetic capacity in *ex vivo* activated CD8^+^ T cells. A-B. Exponentially dividing **(A)** mouse embryonic fibroblasts and **(B)** HeLa cells were treated with Vehicle (Veh) or 5 µM ATRi for 6 h, inhibitor was washed out, and cells were allowed to recover for 18 h. During the final 2 h of the recovery period, mitochondria were stained with MitoView Fix 640, and then cells were fixed for microscopy. Shown are representative fields with scale bars, and **(B)** enlarged fields of individual HeLa cells. **C-H.** CD8^+^ T cells were isolated and activated as described in Figure 1. **C.** Activated CD8^+^ T cells were treated with Veh or 5 µM ATRi for 6 h, inhibitor was washed out, and cells were allowed to recover for 18 h recovery prior to staining with MitoTracker Green (MTG). Shown are representative histograms of MTG fluorescence intensity and quantitation of MTG median fluorescence intensity (MFI), normalized to the mean of Veh controls. Data from one experiment with three separately activated replicates (from two pooled spleens) per group. Mean ± SD bars shown. **p<0.01 by unpaired, two-tailed Welch’s t-test. **D-H.** Activated CD8^+^ T cells were treated with Veh or 5 µM ATRi for 6 h either prior to an 18 h recovery period or following an 18 h expansion period, as indicated. At 48 h hours, metabolic flux analyses were performed using the Seahorse XFe96 Analyzer. Data from one experiment. Each data point or bar represent the mean of eight technical replicates (two replicates for ATRi in panels D and G due to low number of cells), with SEM bars shown. **D-E.** Oxygen consumption rates (OCR, pmol/min) over time for each of the ATRi treatment schedules. **F.** Quantitation of spare respiratory capacity, calculated as maximal OCR - basal OCR. ****p<0.001, ****p<0.0001 by one-way ANOVA with Šidák ‘s multiple comparisons, with no comparisons across treatment schedules. **G-H.** Extracellular acidification rate (ECAR, mpH/min) over time for each of the ATRi treatment schedules.

Clonally expanding CD8^+^ T cells must undergo coordinated and tightly regulated mitochondrial biogenesis, fission, and fusion as the cells rapidly cycle and divide (42-44). We therefore questioned whether temporal dysregulation of fission altered mitochondrial mass in ATRi-treated CD8^+^ T cells. We isolated CD8^+^ T cells from wildtype C57BL/6 mice, activated the cells for 24 h *ex vivo* using anti-CD3 and anti-CD28 antibodies and IL-2, treated the activated CD8^+^ T cells with vehicle or 5 µM ATRi for 6 h, and then washed out inhibitor and allowed cells to recover in IL-2 for an additional 18 h. ATRi significantly reduced mitochondrial mass compared to vehicle control, as measured by MitoTracker Green FM (MTG) staining and flow cytometry (Fig 2C). We performed additional experiments in which we activated CD8^+^ T cells in the same manner, expanded the cells in IL-2 for an additional 24 h or 48 h, treated with ATRi for 6 h, and either immediately stained cells with MTG (Supplemental Fig S1B) or washed out the inhibitor and allowed cells to recover in IL-2 for an additional 18 h (Supplemental Fig S1C). In both cases, ATRi reduced mitochondrial mass in the clonally expanding CD8^+^ T cells (Supplemental Fig S1B-C). Therefore, ATRi increases premature CDK1-dependent DRP1 activity, dysregulates mitochondrial fission, and reduces mitochondrial mass in clonally expanding CD8^+^ T cells.

Since mitochondrial mass was reduced in ATRi-treated cells, we hypothesized that mitochondrial function may also be compromised in ATRi-treated cells. To test this, we used a Seahorse XF Analyzer to assess oxygen consumption rate (OCR), a measure of mitochondrial respiration, and extracellular acidification rate (ECAR), a measure of glycolytic activity, in activated CD8^+^ T cells (isolated from wildtype C57BL/6 mice, activated for 24 h) that were treated with vehicle or ATRi for 6 h. The CD8^+^ T cells were either treated immediately after activation and then allowed to recover in IL-2, without ATRi, for 18 h (Fig 2D) or were expanded in IL-2 for 18 h post-activation and then treated for the final 6 h of the 48-h period (Fig 2E), prior to Seashore analyses at 48 h. Regardless of when the activated CD8^+^ T cells were treated, ATRi reduced OCR by approximately 34-36% (Fig 2D-E) and spare respiratory capacity (maximal OCR minus basal OCR) by approximately 37-47% (Fig 2F) in clonally expanding CD8^+^ T cells. ATRi also reduced ECAR in activated CD8^+^ T cells, regardless of the treatment timing (Fig 2G-H), indicating that overall bioenergetic demand was reduced by ATRi. In the case of clonally expanding CD8^+^ T cells treated during the final 6 h, we also assessed whether addition of exogenous thymidine (dT) could rescue bioenergetic activity of ATRi-treated cells, as we previously reported that dT rescues ATRi-induced cell death in activated CD8^+^ T cells (7). Thymidine did not rescue OCR, ECAR, or spare respiratory capacity in ATRi-treated, clonally expanding CD8^+^ T cells (Supplemental Fig S1D-F), indicating that the reduced bioenergetic capacity after ATRi is not due to cell death.

### ATR signaling is essential to establish CD8^+^ T cell memory

Mitochondrial function in newly activated CD8^+^ T cells is critical for their expansion and differentiation, effector function, and the development of long-lived memory cells (42, 44-46). Since we observed that ATR kinase inhibitors impacted the mitochondria of proliferating CD8^+^ T cells activated *ex vivo*, we reasoned that disruption of mitochondrial function in clonally expanding CD8^+^ T cells may have consequences on CD8^+^ T cell memory. To determine the impact of ATRi on CD8^+^ T cell memory *in vivo*, we infected C57BL/6 mice with LCMV Armstrong and then treated mice with vehicle or ATRi on days 5, 6, and 7, which is within the period of peak CD8^+^ T cell clonal expansion following LCMV Armstrong infection (47). We then re-challenged mice with viral antigen at 6 weeks post infection by injecting them in both flanks with LCMV glycoprotein-expressing B16 melanoma cells (B16-GP). Tumor growth and survival (time to reach tumor endpoint) were monitored from 7 weeks post infection. A schematic of this experimental design is shown in Figure 3A.

**Figure 3.**
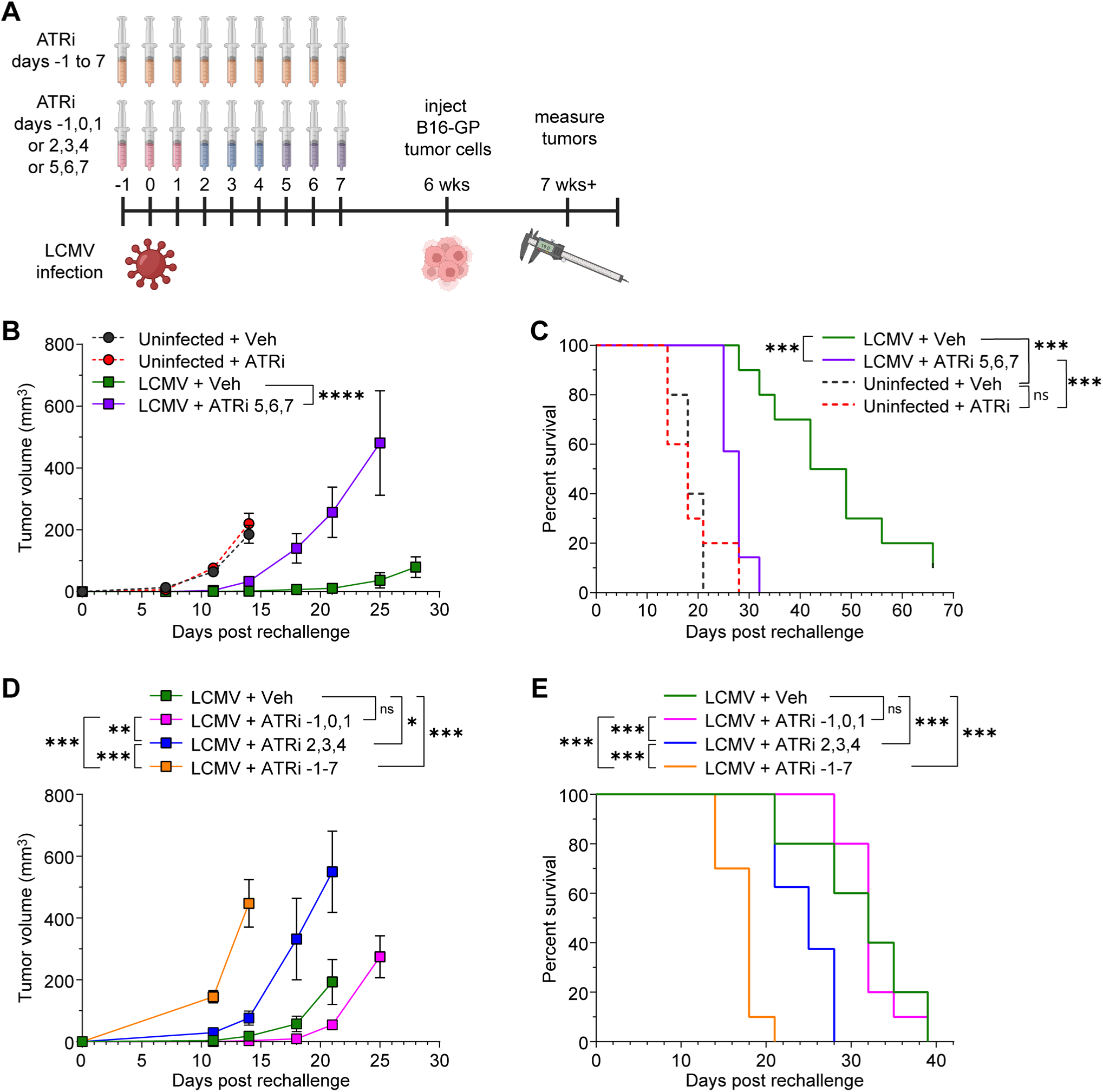
ATR signaling is essential to establish CD8^+^ T cell memory. **A.** Schematic of the experimental timeline. Mice were infected with LCMV Armstrong on day 0, treated as indicated, and then injected on both flanks with LCMV glycoprotein-expressing B16 (B16-GP) tumor cells at 6 weeks post infection. Tumor growth was monitored starting >7 weeks post infection. Uninfected controls groups were treated, B16-GP challenged, and monitored in the same ways. **B-C.** Mice were treated with Vehicle (Veh) or 75 mg/kg ATRi on days 5, 6, and 7 post-infection. **B.** B16-GP tumor growth curves showing mean tumor volume (±SEM). Curves end once more than one mouse was euthanized within a given group. ****p<0.0001 by mixed effects model comparing LCMV-infected groups. **C.** Survival curves for each LCMV-infected treatment group and corresponding uninfected control groups. The survival endpoint was reached when tumor length exceeded 15 mm or tumors ulcerated. ***p<0.001 by log-rank test with Holm-Šidák adjustment for multiple comparisons (all comparisons shown). **B-C.** Data from one experiment with n = 7 mice (14 tumors) for LCMV + ATRi 5,6,7 and n = 10 mice (20 tumors) for all other groups. **D-E.** Mice were treated with Veh (days 2, 3, 4) or with different schedules of 75 mg/kg ATRi as indicated (days -1, 0, 1; days 2, 3, 4; or days -1 to 7). **D.** B16-GP tumor growth curves showing mean tumor volume (±SEM). Curves end once more than one mouse is euthanized within a given group. *p<0.05, **p<0.01, ***p<0.001 by mixed effects model with Holm-Šidák adjustment for multiple comparisons (all LCMV-infected groups). **E.** Survival curves for each LCMV-infected treatment group. The survival endpoint was reached when tumor length exceeded 15 mm or tumors ulcerated. ***p<0.001 by log-rank test with Holm-Šidák adjustment for multiple comparisons (all groups). **D.** Individual tumor growth curves. **D-E.** Data from one experiment with n = 8 mice (16 tumors) for LCMV + ATRi 2,3,4 and n = 10 mice (20 tumors) for all other groups.

B16-GP tumors grew rapidly in uninfected control mice, causing pressure necrosis and/or tumor ulceration that necessitated euthanasia in each group as early as day 14 post-rechallenge (Supplemental Fig S2A). As expected, LCMV Armstrong infection suppressed B16-GP tumor growth (Fig 3B and Supplemental Fig S2A) and prolonged the survival of tumor bearing mice compared to uninfected controls (Fig 3C). Administration of ATRi on days 5, 6, and 7 post-infection attenuated the suppression of B16-GP tumor growth (Fig 3B and Supplemental Fig S2A) and shortened survival compared to LCMV-infected vehicle control mice (median 28 days versus 45.5 days) (Fig 3C). We repeated this experiment injecting B16-GP tumors in a single flank and obtained similar results (Supplemental Fig S2B). These data suggest that ATRi on days 5, 6, and 7 post-LCMV Armstrong infection impairs immune memory.

To determine whether the ATRi treatment schedule post-LCMV Armstrong infection impacted immune memory, we treated mice with ATRi on days -1, 0, and 1, during the initial antigen exposure, days 2, 3, and 4, prior to the period of maximum effector expansion, or days -1 through 7 (Figure 3A). We added an LCMV-infected vehicle control group injected with parental B16 cells (i.e. no LCMV glycoprotein expression) to demonstrate the specificity of our system (Supplemental Fig S1E).

Treatment with ATRi on days -1, 0, and 1 had no significant impact on tumor growth or survival following rechallenge (Fig 3D-E), suggesting that ATR signaling is not required for priming of CD8^+^ T cell effector responses. ATRi on days 2, 3, and 4 post-infection attenuated the suppression of B16-GP tumor growth by prior LCMV Armstrong infection (Fig 3D and Supplemental Fig S2C) and shortened survival compared to LCMV-infected vehicle control mice (median 25 days versus 32 days) (Fig 3E). ATRi on days -1 through 7 post-infection profoundly limited tumor control (Fig 3D and Supplemental Fig S2C) and significantly reduced median survival to 18 days (Fig 3E). Together, these data suggest that exposure of clonally expanding CD8^+^ T cells to ATRi has lasting impacts on CD8^+^ T cell memory, and that ATR signaling during CD8^+^ T cell clonal expansion is essential for the proper development of CD8^+^ T cell memory.

### ATR signaling is essential for CD8^+^ T cell clonal expansion after viral infection

Next, we investigated the impact of ATRi on the effector CD8^+^ T cell response in splenocytes at day 8 post-LCMV Armstrong infection (Fig 4A). ATRi administration on days 2, 3, and 4, or 5, 6, and 7, or -1-7 post-LCMV Armstrong infection, during CD8^+^ T cell clonal expansion, significantly reduced splenic CD8^+^ T cells, while ATRi on days -1, 0, and 1, prior to clonal expansion, had no impact on CD8^+^ T cell number (Fig 4B). We examined CD44 and CD26L expression on splenic CD8^+^ T cells to determine their activation and maturation phenotypes (Fig 4C). Mirroring the impact of ATRi on the abundance of splenic CD8^+^ T cells, the proportions of effector/effector memory CD8^+^ T cells (T_EM,_ CD44^+^CD62L^neg^) were significantly reduced in all groups receiving ATRi during CD8^+^ T cell clonal expansion (Fig 4C-E). Concomitantly, the proportions of central memory (T_CM,_ CD44^+^CD62L^+^) and naïve (T_N,_ CD44^neg^CD62L^+^) CD8^+^ T cells were significantly increased in these mice (Fig 4C-E). The impact of ATRi on days 5, 6, and 7, and days -1-7 on CD8^+^ T cell activation and maturation was more profound than ATRi on days 2, 3, and 4, while ATRi on days -1, 0, and 1 had no impact on these subsets (Fig 4C-E).

**Figure 4.**
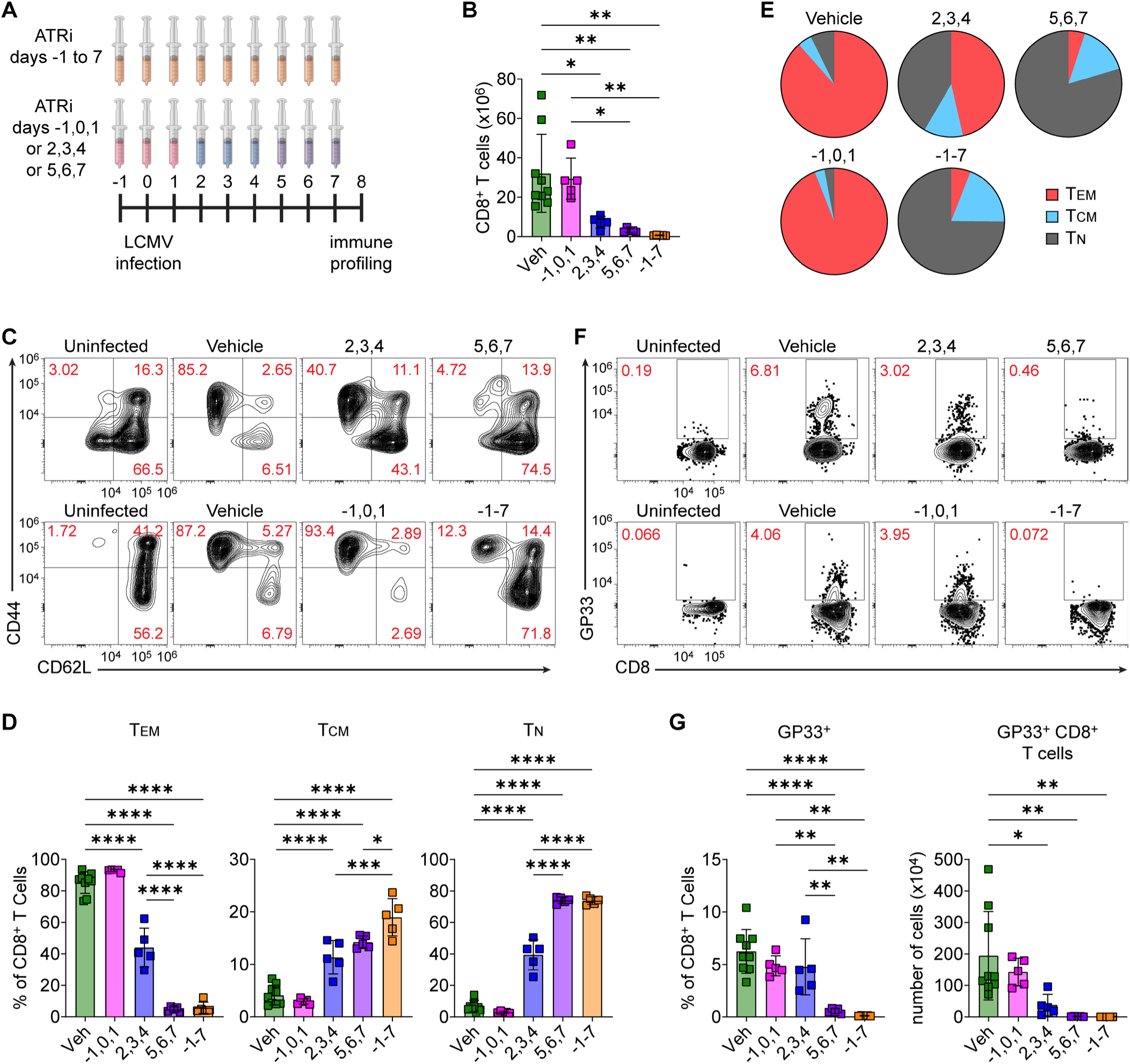
ATR signaling is essential for virus-specific CD8^+^ T cell clonal expansion. **A.** Schematic of the experimental timeline. Mice were infected with LCMV Armstrong on day 0 and were treated daily with Vehicle (Veh, days 2, 3, 4 or days 5, 6, 7) or with different schedules of 75 mg/kg ATRi as indicated (days -1, 0, 1; days 2, 3, 4; or days -1 to 7). Splenocytes were immune profiled by spectral flow cytomety on day 8. **B.** Quantitation of the number of CD8^+^ T cells per spleen sample. **C.** Representative contour plots showing CD44 and CD62L expression on CD8^+^ T cells from uninfected control, Veh-treated, and ATRi-treated mice (treatment days indicated), with the top row from one experiment and the bottom row from a second experiment. **D.** Quantification of CD44^+^CD62L^neg^ effector/effector memory (TEM), CD44^+^CD62L^+^ central memory (TCM), and CD44^neg^CD62L^+^ naïve (TN) CD8^+^ T cells, as percentages of total CD8^+^ T cells. **E.** Pie charts showing the mean fractions of CD8^+^ TEM, TCM, and TN per group. **F.** Representative contour plots showing GP33^+^ CD8^+^ T cells from uninfected control, Veh-treated, and ATRi-treated mice (treatment days indicated), with the top row from one experiment and the bottom row from a second experiment. **G.** Quantitation of GP33^+^ CD8^+^ T cells, both as a percentage of CD8^+^ T cells and as the number of GP33^+^ CD8^+^ T cells per spleen sample. **B-G.** Data combined from two experiments, each with n = 5 mice per group per experiment, for a total n = 10 Veh and n = 5 for each ATRi treatment group. One uninfected control was included in each experiment. **B, D, G**. Individual data points with mean ± SD bars shown. *p<0.05, **p<0.01, ***p<0.001 ****p<0.0001 by one-way ANOVA with Tukey’s multiple comparisons test. **B, G.** All statistically significant comparisons shown. **D.** Statistically significant comparisons not shown for -1,0,1 versus the remaining ATRi groups (2,3,4 or 5,6,7 or -1 to 7), but are identical to Veh versus those ATRi groups. All other statistically significant comparisons are shown.

We examined KLRG1 and CD127 expression in the total CD8^+^ T cell population (Supplemental Fig S3A), rather than in T_EM_ specifically, given the limited abundance of T_EM_ in the mice that received ATRi post-LCMV infection on days 5, 6, and 7 or on days -1-7. KLRG1^+^CD127^lo^ CD8^+^ T cells generally represent short-lived effector cells, while KLRG1^neg^CD127^hi^ CD8^+^ T cells generally represent the memory precursor compartment, although these phenotypic distinctions are not absolute, and KLRG1^+^CD127^lo^ long-lived effector cells have been described (38, 48, 49). The distribution of these phenotypes mirrored that observed with the T_EM_ and T_CM_ subsets for all treatment groups. Dosing animals with ATRi on days 5, 6, and 7 or days -1-7 post-LCMV Armstrong nearly eliminated KLRG1^+^CD127^lo^ cells and proportionally increased KLRG1^neg^CD127^hi^ cells. ATRi on days 2, 3, and 4 had a less substantial impact on these subsets (Supplemental Fig S3B), likely due to partial recovery of CD8^+^ T cell expansion and maturation recovery following cessation of ATRi (23, 25). ATRi on days -1, 0, and 1 was again indistinguishable from the vehicle-treated group (Supplemental Fig S3B).

LCMV infection of C57BL/6 mice triggers expansion of antigen-specific T cells recognizing a wide range of epitopes. One of the three immunodominant epitopes, GP33, is an MHC class I H2-D^b^ restricted peptide derived from residues 33-41 of the LCMV glycoprotein (GP). We utilized MHC class I tetramer loaded with GP33 peptide to assess GP33-specific CD8^+^ T cell responses in LCMV-infected mice treated with the vehicle or the ATRi schedules described above (Fig 4F). Splenic GP33^+^ CD8^+^ T cells were nearly eliminated by ATRi treatment on days 5, 6, and 7, or days -1-7 (Fig 4F-G). While ATRi on days 2, 3, and 4 did not significantly alter the frequency of GP33^+^ cells within the CD8^+^ T cell pool, it did significantly reduce the overall abundance of splenic GP33^+^ CD8^+^ T cells (Fig 4F-G). ATRi on days -1, 0, and 1 did not impact the frequency or abundance of splenic GP33^+^ CD8^+^ T cells (Fig 4G).

Of the three-day schedules, ATRi on days 5, 6, and 7 had the most profound effects on CD8^+^ T cell clonal expansion, and on activation and maturation phenotypes of the overall and GP33-specific CD8^+^ T cell pools. We therefore studied the effect of treatment with ATRi on days 5, 6, and 7 after infection in more detail.

### Cessation of ATRi triggers a delayed effector CD8^+^ T cell response

We hypothesized that ATRi-treated mice mount an effector CD8^+^ T cell response after cessation of treatment. To test this, we immune profiled splenocytes on day 18 in mice treated with ATRi on days 5, 6, and 7 post-LCMV Armstrong infection (Fig 5A). Day 18 falls within the T cell contraction phase post-LCMV infection (37, 47). ATRi on days 5, 6, and 7 did not impact the abundance of splenic CD8^+^ T cells at day 18 (Fig 5B), suggesting that CD8^+^ T cells proliferated after cessation of ATRi. Furthermore, proliferating (Ki67^+^) CD8^+^ T cells were, on average, increased in ATRi-treated mice, although this did not reach statistical significance due to large variation in the ATRi group (Supplemental Fig S4A). We examined CD44 and CD26L expression on splenic CD8^+^ T cells (Supplemental Fig S4B) and found that, while the CD8^+^ T_EM_ subset had expanded substantially in ATRi-treated mice, it remained lower than in vehicle-treated mice (Fig 5C). Conversely, the T_N_ compartment was proportionally increased in ATRi-treated animals (Supplemental Fig S4C).

**Figure 5.**
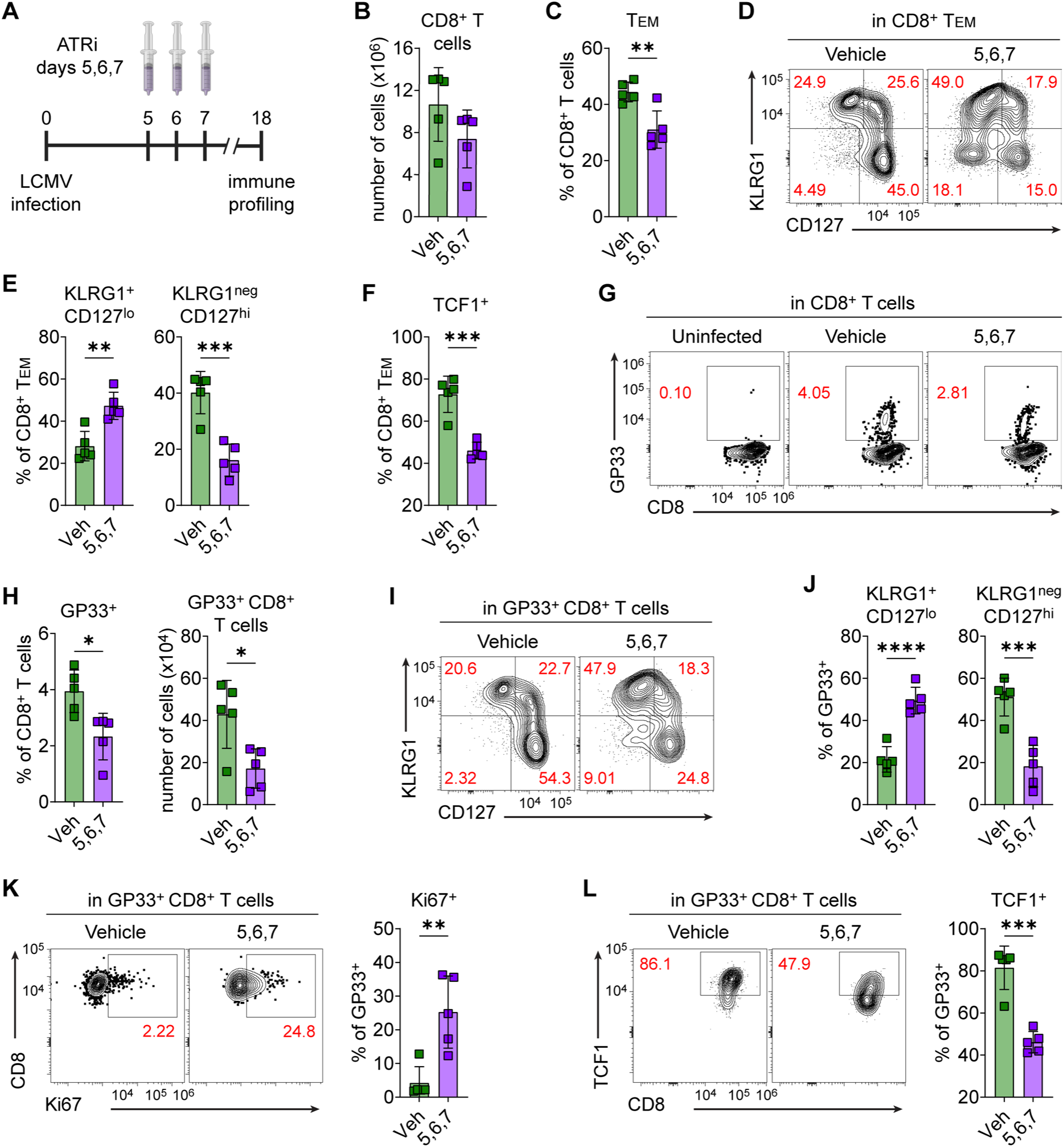
Cessation of ATRi triggers a delayed effector CD8^+^ T cell response. **A.** Schematic of the experimental timeline. Mice were infected with LCMV Armstrong on day 0 and were treated daily with Vehicle (Veh) or 75 mg/kg ATRi (days 5, 6, 7). Splenocytes were immune profiled by spectral flow cytomety on day 18. **B.** Quantitation of the number of CD8^+^ T cells per spleen sample. **C.** Quantitation of CD8^+^ TEM as percentages of total CD8^+^ T cells. **D.** Representative contour plots showing KLRG1 and CD127 expression on CD8^+^ TEM from Veh-treated and ATRi-treated mice. **E.** Quantification of KLRG1^+^CD127^lo^ and KLRG1^neg^CD127^hi^ CD8^+^ TEM, as percentages of total CD8^+^ TEM. **F.** Quantification of TCF1^+^ CD8^+^ TEM, as a percentage of total CD8^+^ TEM. **G.** Representative contour plots showing GP33^+^ CD8^+^ T cells from uninfected control, Veh-treated, and ATRi-treated mice. **H.** Quantitation of GP33^+^ CD8^+^ T cells, both as a percentage of CD8^+^ T cells and as the number of GP33^+^ CD8^+^ T cells per spleen sample. **I.** Representative contour plots showing KLRG1 and CD127 expression on GP33^+^ CD8^+^ T cells from Veh-treated and ATRi-treated mice. **J.** Quantification of KLRG1^+^CD127^lo^ and KLRG1^neg^CD127^hi^ GP33^+^ CD8^+^ T cells, as percentages of total GP33^+^ CD8^+^ T cells. **K-L.** Representative contour plots showing **(K)** Ki67 or **(L)** TCF1 expression in GP33^+^ CD8^+^ T cells from Veh-treated and ATRi-treated mice, and quantification of **(K)** Ki67^+^ or **(L)** TCF1^+^ GP33^+^ CD8^+^ T cells, as percentages of total GP33^+^ CD8^+^ T cells. **B-L.** Data from one experiment with n = 5 mice per group and including one uninfected control. **B-C, E-F, H, J-L.** Individual data points with mean ± SD bars shown. *p<0.05, **p<0.01, ***p<0.001 ****p<0.0001 by unpaired, two-tailed Welch’s t-test.

KLRG1 and CD127 expression of the CD8^+^ T_EM_ population revealed that most of the T_EM_ pool in ATRi-treated mice exhibited a short-lived effector phenotype (KLRG1^+^CD127^lo^), whereas the T_EM_ of vehicle-treated mice predominantly exhibited a memory precursor (KLRG1^neg^CD127^hi^) phenotype (Fig 5D-E). ATRi-treated mice had significantly more KLRG1^neg^CD127^lo^ double-negative cells in their CD8^+^ T_EM_ pools (Supplemental Fig S4D). Since these double-negative cells generally represent an early effector differentiation state (50-52), an increase in this population at day 18 is consistent with a recovering, but not yet fully developed, effector CD8^+^ T cell response in ATRi-treated mice. Conversely, The CD8^+^ T_EM_ of vehicle-treated mice had significantly more KLRG1^+^CD127^hi^ double-positive cells (Supplemental Fig S4D), which generally represents more differentiated memory precursors and long-lived effector cells (38, 48, 53).

CD8^+^ T_EM_ had greater proportions of memory and memory-like cells in vehicle-treated mice, while this population skewed toward early effector and short-lived effector cells in ATRi-treated mice. Furthermore, the CD8^+^ T_EM_ had a significantly greater frequency of cells expressing the “stemness” associated transcription factor TCF1 in vehicle-treated mice (Fig 5F). While the frequency of TCF1^+^ cells also increased in total CD8^+^ T cell pools in vehicle-treated mice (Supplemental Fig S4E), we observed no differences in the frequency of TCF1^+^ cells within the CD8^+^ T_CM_ of vehicle-treated and ATRi-treated mice (Supplemental Fig S4F). Therefore, ATRi decreases the TCF1^+^ stem-like, multipotent, and highly proliferative cells in the CD8^+^ T_EM_ population, which are critical for replenishing effector CD8^+^ T cell pools in infected and tumor-bearing mice (54-58).

Next, we examined the same populations within the defined GP33-specific CD8^+^ T cell pools (Fig 5G). The frequency and abundance of GP33^+^ CD8^+^ T cells remained significantly lower in ATRi-treated mice by day 18 but had expanded considerably since cessation of ATRi treatment (Fig 5G-H). Examination of CD44 and CD62L expression in the GP33^+^ population revealed that the frequency of T_CM_ phenotype cells in ATRi-treated mice was approximately half of that found in vehicle-treated mice (Supplemental Fig S4G-H). ATRi-treated mice had significantly increased frequencies of short-lived effector (KLRG1^+^CD127^lo^) cells within their GP33^+^ pools, while vehicle-treated mice had significantly higher frequencies of memory precursor (KLRG1^neg^CD127^hi^) cells (Fig 5I-J). On average, KLRG1^neg^CD127^lo^ (double-negative) GP33^+^ CD8^+^ T cells were present at higher frequency in ATRi-treated mice, while KLRG1^+^CD127^hi^ double-positive GP33^+^ CD8^+^ T cells were higher in vehicle-treated mice, but these differences did not reach statistical significance due to large variation in ATRi-treated mice (Supplemental Fig S4I).

At day 18, the GP33^+^ CD8^+^ T cell pools of ATRi-treated mice had increased frequencies of actively proliferating (Ki67^+^) cells (Fig 5K), reduced frequencies of TCF1-expressing cells (Fig 5L), and increased expression of the effector T cell-associated transcription factor T-bet (based on median fluorescence intensity, MFI) in the T-bet^+^ pools (Supplemental Fig S4J). Taken together, these data suggest that cessation of ATRi treatment triggers an effector CD8^+^ T cell response with delayed kinetics and with greater bias toward short-lived effector and less-differentiated early effector cells, rather than effector memory precursor and more-differentiated effector memory cells in LCMV Armstrong infected mice.

### ATRi during CD8^+^ T cell clonal expansion permanently alters memory CD8^+^ T cell phenotypes

Next, we interrogated the impact of days 5, 6, and 7 ATRi treatment on virus-specific memory CD8^+^ T cells by immune profiling splenocytes at 7 weeks post LCMV infection (Fig 6A). At this late time point, we did not observe any impact of prior ATRi treatment on the total number of splenic CD8^+^ T cells (Fig 6B), but ATRi did modestly reduce the CD8^+^ fraction in CD45^+^ splenocytes (Supplemental Fig S5A). Interestingly, there was elevated CD8^+^ T cell proliferation (Ki67^+^) in ATRi-treated mice compared to vehicle-treated mice, even at this late 7-week post-infection timepoint (Supplemental Fig S5B). ATRi also caused a small, but statistically significant, reduction in the frequency of T_CM_ in total CD8^+^ T cells (Supplemental Fig S5C-D). We did not observe any significant differences in the KLRG1/CD127 phenotypic subsets (Supplemental Fig S5E-F). However, ATRi caused significant reductions in the frequency of stem-like TCF1^+^ cells in the total CD8^+^ T cell and, particularly, in the CD8^+^ T_EM_ pools (Supplemental Fig S5G-H). ATRi also caused modestly increased frequencies of T-bet^+^ cells and T-bet expression (based on MFI) within the T-bet^+^ pools of CD8^+^ T cells (Supplemental Fig S5I-J), signifying a more effector-like memory pool.

**Figure 6.**
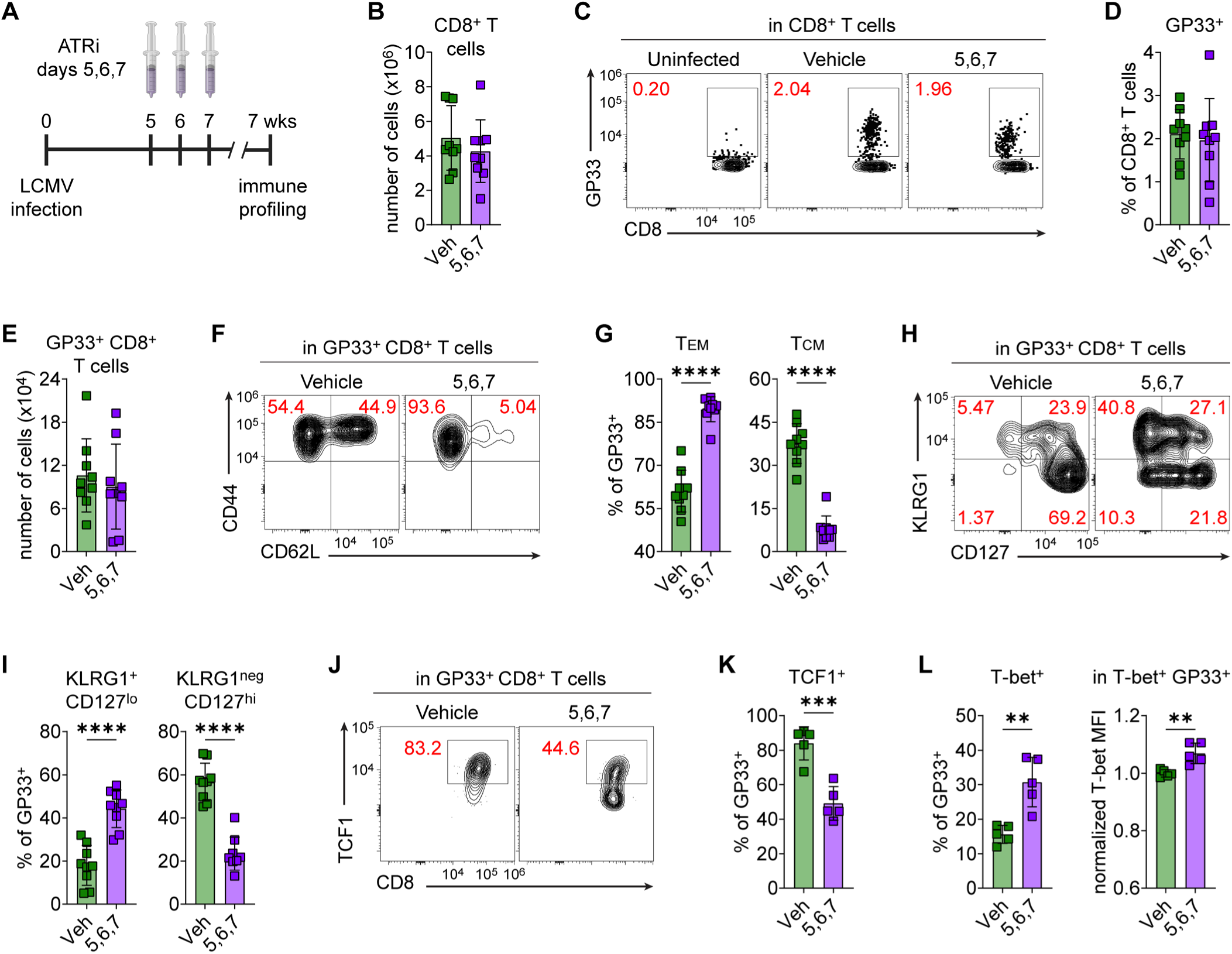
ATRi during CD8^+^ T cell clonal expansion permanently alters memory CD8^+^ T cell phenotypes. **A.** Schematic of the experimental timeline. Mice were infected with LCMV Armstrong on day 0 and were treated daily with Vehicle (Veh) or 75 mg/kg ATRi (days 5, 6, 7). Splenocytes were immune profiled by spectral flow cytometry at 7 weeks post infection. **B.** Quantitation of the number of CD8^+^ T cells per spleen sample. **C.** Representative contour plots showing GP33^+^ CD8^+^ T cells from uninfected control, Veh-treated, and ATRi-treated mice. **D-E.** Quantitation of GP33^+^ CD8^+^ T cells, both as **(D)** a percentage of CD8^+^ T cells and as **(E)** the number of GP33^+^ CD8^+^ T cells per spleen sample. **F.** Representative contour plots showing CD44 and CD62L expression on GP33^+^ CD8^+^ T cells from Veh-treated and ATRi-treated mice. **G.** Quantification of GP33^+^ CD8^+^ TEM and TCM, as percentages of total GP33^+^ CD8^+^ T cells. **H.** Representative contour plots showing KLRG1 and CD127 expression on GP33^+^ CD8^+^ T cells from Veh-treated and ATRi-treated mice. **I.** Quantification of KLRG1^+^CD127^lo^ and KLRG1^neg^CD127^hi^ GP33^+^ CD8^+^ T cells, as percentages of total GP33^+^ CD8^+^ T cells. **J.** Representative contour plots showing TCF1 expression in GP33^+^ CD8^+^ T cells from Veh-treated and ATRi-treated mice. **K.** Quantification of TCF1^+^ GP33^+^ CD8^+^ T cells, as percentages of total GP33^+^ CD8^+^ T cells. **L.** Quantification of T-bet^+^ GP33^+^ CD8^+^ T cells, as a percentage of total GP33^+^ CD8^+^ T cells, and quantitation of T-bet median fluorescence intensity (MFI), normalized to the mean of Veh-treated mice, in T-bet^+^ GP33^+^ CD8^+^ T cells. **B-I.** Data combined from two independent experiments, each with n = 4-5 mice per group, for total n = 9 mice per group. Each experiment included one uninfected control. **J-L.** Data from one of the two above experiments, with n = 5 mice per group. **B, D, E, G, I, K-L.** Individual data points with mean ± SD bars shown. *p<0.05, **p<0.01, ****p<0.0001 by unpaired, two-tailed Welch’s t-test.

GP33^+^ memory pools stabilized in ATRi-treated mice by 7 weeks post LMCV-infection, as we observed no differences between ATRi- and vehicle-treated mice in the frequency or abundance of splenic GP33^+^ CD8^+^ T cells (Fig 6C-E), or in the frequency of proliferating (Ki67^+^) GP33^+^ CD8^+^ T cells (Supplemental Fig S5K). Strikingly, ATRi on days 5, 6, and 7 resulted in a profound and lasting shift in the memory subsets within the GP33^+^ pools (Fig 6F). ATRi-treated mice had a significantly higher proportion of GP33^+^ T_EM_ and significantly reduced GP33^+^ T_CM_ (Fig 6G). In addition, ATRi-treated mice had significantly higher frequencies of KLRG1^+^CD127^lo^ cells and significantly reduced frequencies of memory precursor KLRG1^neg^CD127^hi^ cells within their GP33^+^ pools (Fig 6H-I), suggesting an overall more effector-like memory pool. Consistent with this premise, the GP33^+^ CD8^+^ T cell pools of ATRi-treated mice had reduced frequencies of TCF1-expressing cells (Fig 6J-K) and increased frequencies of T-bet-expressing cells with elevated T-bet expression (Fig 6L and Supplemental Fig S5L). Taken together, ATRi on days 5, 6, and 7 results in permanent changes in the GP33-specific CD8^+^ T cell memory pools, with stark shifts toward effector-like and away from stem-like memory cells.

### ATRi during CD8^+^ T cell clonal expansion reduces the proliferative capacity of the memory CD8^+^ T cell pool

ATRi on days 5, 6, and 7 post-LCMV Armstrong infection impairs CD8^+^ T cell memory recall, and this is associated with lasting changes in the GP33^+^ CD8^+^ T cell memory compartment, including reductions in central memory and stem-like (TCF1^+^) memory cells and shifts toward effector-like (KLRG1^+^, T-bet^+^) memory cells. Effector memory cells are known to possess more immediate cytokine function and reduced proliferative capacity compared to central memory cells, making effector memory cells more suited to on-the-spot surveillance, while central memory cells are more suited to generate full-fledged secondary effector responses (59-63). Therefore, we hypothesized that the GP33^+^ memory compartments of ATRi-treated mice would exhibit increased cytokine producing capacities and decreased proliferative potential. To test this, we stimulated splenocytes from ATRi-treated and vehicle-treated mice at 12 weeks post LCMV-infection with GP33 peptide (Fig 7A). We waited until 12 weeks post infection to eliminate any question as to whether the GP33^+^ memory pools had entirely stabilized.

**Figure 7.**
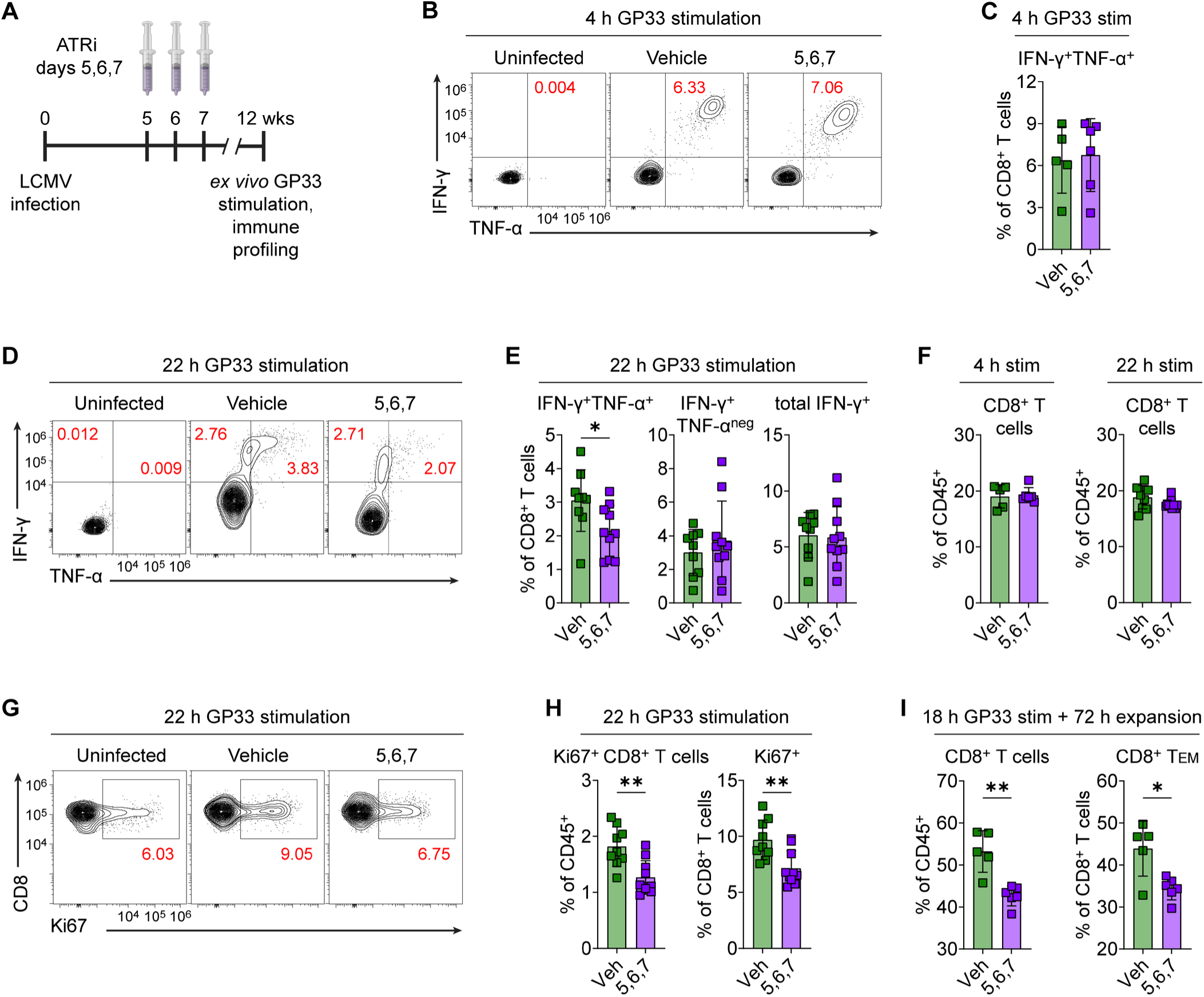
ATRi during CD8^+^ T cell clonal expansion reduces the proliferative capacity of the memory CD8^+^ T cell pool. **A.** Schematic of the experimental timeline. Mice were infected with LCMV Armstrong on day 0 and were treated daily with Vehicle (Veh) or 75 mg/kg ATRi (days 5, 6, 7). Splenocytes were stimulated *ex vivo* with GP33 peptide (for the indicated durations), and then immune profiled by spectral flow cytometry, at 12 weeks post infection. **B-C.** Splenocytes were stimulated with 100 ng/mL GP33 peptide + 25 U/mL IL-2 + protein transport inhibitors for 4 h. **B.** Representative contour plots showing IFN-γ and TNF-α expression in CD8^+^ T cells from uninfected, Veh-treated, and ATRi-treated mice. **C.** Quantitation of IFN-γ^+^TNF-α^+^ CD8^+^ T cells, as a percentage of total CD8^+^ T cells. **D-E.** Splenocytes were stimulated with 100 ng/mL GP33 peptide + 25 U/mL IL-2 for 22 h with protein transport inhibitors added for the final 4 h. **D.** Representative contour plots showing IFN-γ and TNF-α expression in CD8^+^ T cells from uninfected, Veh-treated, and ATRi-treated mice. **E.** Quantitation of IFN-γ^+^TNF-α^+^, IFN-γ^+^TNF-α^neg^, and total IFN-γ^+^ (TNF-α^+^ or TNF-α^neg^) CD8^+^ T cells, as percentages of total CD8^+^ T cells. **F.** Quantitation of CD8^+^ T cells, as percentages of CD45^+^ immune cells, following GP33 stimulation for 4 h (left, same splenocytes as in panels **B-C**) or 22 h (right, same splenocytes as in panels **D-E**). **G-H.** The same splenocytes as in panels **B-C** that were stimulated with GP33 for 22 h. **G.** Representative contour plots showing Ki67 expression in CD8^+^ T cells from uninfected, Veh-treated, and ATRi-treated mice. **H.** Quantitation of Ki67^+^ CD8^+^ T cells, both as a percentage of CD45^+^ immune cells and as a percentage of total CD8^+^ T cells. **I.** Splenocytes were stimulated with 100 ng/mL GP33 peptide + 50 U/mL IL-2 for 18 h and were then expanded in 50 U/mL IL-2 for 72 h. Shown are quantitation of CD8^+^ T cells (as a percentage of CD45^+^ immune cells) and CD8^+^ TEM (as a percentage of total CD8^+^ T cells). **B-C, F (left), I.** Data from one experiment with n = 5 (Veh) or 6 (ATRi) mice. **D-E, F (right), G-H.** Data combined from two independent experiments with n = 4-6 mice per group for total n = 9 (Veh) or 10 (ATRi) mice. **C, E-F, H-I.** Individual data points with mean ± SD bars shown. *p<0.05, **p<0.01 by unpaired, two-tailed Welch’s t-test.

A short (4 h) stimulation with GP33 (100 ng/mL) in the presence of 25 U/mL IL-2 and protein transport inhibitors yielded no between ATRi- and vehicle-treated mice in the IFN-γ and TNF-α producing capacity of the CD8+ T cells (Fig 7B-C). In contrast, when we stimulated splenocytes for 22 h (including protein transport inhibitors for the final 4 h), we found that ATRi-treated mice had a significantly reduced frequency of IFN-γ and TNF-α co-expressing CD8^+^ T cells (Fig 7D-E). Nevertheless, there was no difference between groups in the total frequency of CD8^+^ T cells that expressed IFN-γ (total IFN-γ^+^) independent of TNF-α expression (TNF-α^+^ or TNF-α^neg^) (Fig 7E). We conclude that the effector function of GP33-specific memory CD8^+^ T cells from ATRi-treated mice is weakened compared to vehicle-treated mice but is not entirely impaired.

For both the 4 hour and the 24 h GP33 stimulations, we noted no differences in the abundance of CD8^+^ T cells (as percentages of all CD45^+^ splenocytes stimulated) after stimulation (Fig 7F). Since we seeded equal numbers of splenocytes for stimulation, these data indicate that the starting abundance of CD8^+^ T cells were equivalent between ATRi- and vehicle-treated mice. However, the frequencies of proliferating (Ki67^+^) CD8^+^ T cells, as percentages of both total CD45^+^ immune cells and total CD8^+^ T cells, were significantly reduced in ATRi-treated mice following 22 h GP33 stimulation (Fig 7G-H), suggesting that the GP33-specific CD8^+^ T cell pools of ATRi-treated mice had reduced proliferative capacity compared to vehicle-treated mice.

To confirm this, we used separate dishes of splenocytes, from the same treated mice, that were stimulated with GP33 (100 ng/mL) and IL-2 (50 U/mL) for 18 h, and then re-seeded into new media with IL-2 (50 U/mL) to expand the CD8^+^ T cells for an additional 72 h. Following the 72-h expansion period, splenocytes from ATRi-treated mice had significantly fewer CD8^+^ T cells, as a percentage of total CD45^+^ immune cells, and a significantly reduced frequency of CD8^+^ T_EM_, as a percentage of total CD8^+^ T cells (Fig 7I). These data clearly demonstrate that the GP33-specific memory CD8^+^ T cell pool of ATRi-treated mice had reduced proliferative capacity compared to that of vehicle-treated mice.

### ATRi during CD8^+^ T cell clonal expansion represses mitochondrial genes and reduces mitochondrial mass in memory CD8^+^ T cells

Finally, we examined the underlying transcriptome changes in GP33^+^ memory CD8^+^ T cells that resulted from exposure of the clonally expanding CD8^+^ T cells to ATRi on days 5, 6, and 7. We sorted GP33^+^ memory CD8^+^ T cells from ATRi-treated and vehicle-treated mice at 6 weeks post-infection and performed bulk RNA sequencing (Fig 8A). ATRi on days 5, 6, and 7 post-LCMV Armstrong infection significantly altered expression of 239 genes in the GP33^+^ memory CD8^+^ T cells, with upregulation of 10 genes and downregulation of 229 genes by at least 1.5-fold (Fig 8B). Remarkably, Gene Ontology (GO) analysis of the differentially expressed transcripts revealed significant downregulation of genes involved in biological processes related to mitochondrial function and biosynthesis, and genes encoding mitochondrial and ribosomal (both mitochondrial and cytosolic) components (Fig 8C). Included were numerous nuclear-encoded mitochondrial genes encoding components of Complex I (*Ndufa11*, *Ndufa13, Ndufaf3, Ndufaf8,Ndufb10, Ndufb4c, Ndufb6, Ndufb7, Ndufs5, Ndufs6*), Complex III (*Uqcr11, Uqcrq*), Complex IV (*Cox14*, *Cox17*, *Cox5b*, *Cox6b1*, *Cox8a*), the mitoribosome (*Impdh1, Micos13, Mrpl14, Mrpl54, Mrps21, Mrps24, Mrps36, Mrps5*), and the inner and outer membrane translocases (*Timm10b* and *Tomm6*, respectively), as well as one mitochondrially-encoded gene (*mt-Rnr1*) that encodes the mitochondrial 12S rRNA (Fig 8D). Multiple cytosolic ribosomal protein-encoding genes (*Mrpl14, Mrpl54*, *Mrps21*, *Mrps24*, *Mrps36*, *Mrps5*) were also downregulated following ATRi treatment (Fig 8D).

**Figure 8.**
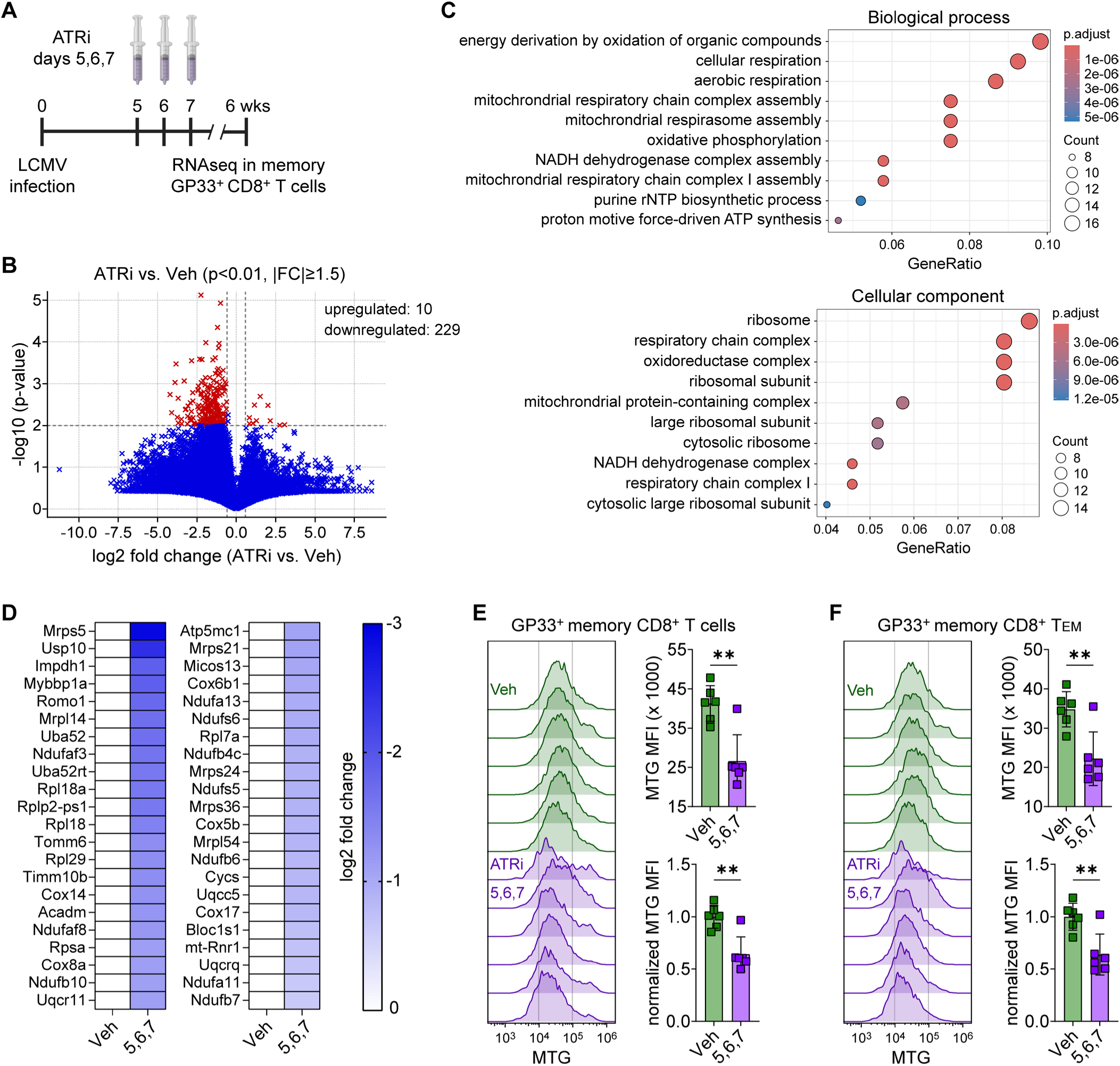
ATRi during CD8^+^ T cell clonal expansion represses mitochondrial genes and reduces mitochondrial mass in memory CD8^+^ T cells. **A.** Schematic of the experimental timeline. Mice were infected with LCMV Armstrong on day 0 and were treated daily with Vehicle (Veh) or 75 mg/kg ATRi (days 5, 6, 7). GP33^+^ memory CD8^+^ T cells were sorted at 6 weeks post infection for RNA sequencing. **B.** Volcano plot depicting the number of differentially expressed genes that were significantly upregulated or downregulated (≥1.5-fold change in expression, p<0.01) in ATRi-treated versus Veh-treated mice. **C.** Gene Ontology (GO) enrichment analysis showing the top 10 most significantly altered biological processes and cellular components in ATRi-treated versus Veh-treated mice. **D.** Heatmaps showing significantly downregulated genes associated with the biological priocesses and cellular components identified in the GO analysis. Genes are ranked from most to least downregulated. **B-D.** n = 5 mice per group. **E-F.** Mice were treated as shown in Panel A and spleens were harvested at 6 weeks post infection for flow cytometry analyses of MitoTracker Green (MTG) staining in GP33^+^ memory CD8^+^T cells. Shown are MTG fluorescence intensity histograms for each mouse and quantitation of both raw and normalized (to the mean of Veh controls) MTG median fluorescence intensity (MFI) in **(E)** total GP33^+^ memory CD8^+^ T cells and **(F)** the GP33^+^ memory CD8^+^ TEM subpopulation. Data from one experiment with n = 6 mice per group. Individual data points with mean ± SD bars shown. **p<0.01 by unpaired, two-tailed Welch’s t-test.

The gene expression changes we observed suggested that the GP33^+^ memory CD8^+^ T cells from ATRi-treated mice were restricted in numerous proteins involved in mitochondrial function and biogenesis and therefore may exhibit reduced overall mitochondrial mass. To test this, we examined mitochondrial mass in the GP33^+^ memory CD8^+^ T cells from vehicle-treated and ATRi-treated mice at 6 weeks post LCMV infection using MitoTracker Green FM (MTG) staining and flow cytometry. MTG is a fluorescent dye that selectively accumulates in mitochondria independent of mitochondrial membrane potential and is widely used to visualize mitochondrial morphology and mass in live cells.

The GP33^+^ memory CD8^+^ T cells from ATRi-treated mice exhibited significantly reduced MTG staining compared to vehicle-treated, either quantified as overall MFI or MFI normalized to the mean of Vehicle-only controls (Fig 8E). Immune profiling data at 7 weeks post-LCMV Armstrong infection showed that a significantly larger portion of GP33^+^ memory CD8^+^ T cell pool in ATRi-treated mice was comprised of effector memory rather than central memory cells. Effector memory CD8^+^ T cells rely more heavily on glycolysis to meet energetic demands, while central memory cells rely more on oxidative phosphorylation and fatty acid oxidation for their energetic needs (42, 44, 45).

Therefore, we reasoned that the reduced mitochondrial mass (MTG staining) in the GP33^+^ memory CD8^+^ T cell pool from ATRi-treated mice may simply reflect the greater proportion of GP33^+^ CD8^+^ T_EM_ that have reduced mitochondrial demand. However, this was not the case, as overall MTG MFI and normalized MTG MFI were significantly lower in the GP33^+^ CD8^+^ T_EM_ pools of ATRi-treated mice than in the GP33^+^ CD8^+^ T_EM_ pools of Vehicle-treated mice (Fig 8F). Therefore, exposure of clonally expanding, virus-specific CD8^+^ T cells to ATRi on days 5, 6, and 7 post-LCMV Armstrong infection leads to reduced mitochondrial mass in the resultant GP33^+^ memory CD8^+^ T cells.

## DISCUSSION

We show that ATR inhibition induces CDK1-dependent phosphorylation of DRP1 at serine-616, triggering mitochondrial fission. These findings underscore ATR signaling as a core regulator of the unperturbed cell cycle that temporally restricts CDK1 activity to mitosis. ATR and its effector CHK1 constitute a brake on CDK1 activity that operates across multiple substrates through S phase and G2 in unperturbed cells (64). ATR signaling restrains CDK1-dependent phosphorylation of several targets: RRM2, whose phosphorylation triggers its degradation in mitosis (7, 65); FOXM1, whose phosphorylation drives transcription at the G2/M transition (6); RIF1, phosphorylation of which limits origin firing throughout S phase (5, 66); and, as we now show, DRP1, whose phosphorylation induces mitochondrial fission in mitosis. Consequently, ATR kinase inhibitors elicit multiple forms of damage in otherwise unperturbed asynchronous cells including deoxyuridine incorporation at replication forks as a result of an elevated dUTP/TTP ratio, as well as premature transcription, aberrant origin firing, and untimely mitochondrial fission. Replication forks that subsequently stall or collapse can neither be stabilized nor limited by checkpoint signaling as they would be following exposure to other chemotherapies, since ATR is inhibited. These effects of ATRi are particularly pronounced in rapidly dividing CD8^+^ T cells (7).

Our data reveal an exquisitely sensitive temporal requirement for mitochondrial fission during effector expansion in the establishment of CD8^+^ T cell memory. The fate decision between terminal effector and memory precursor cells occurs during a critical window of days 4–8 post-LCMV Armstrong infection, with the subsequent contraction phase (days 9–30) shaping how many memory cells ultimately survive (37, 38). CD8^+^ T memory cells harbor more mitochondria and have a higher reliance on oxidative phosphorylation for greater ATP production capacity than naïve CD8^+^ T cells, and this enhanced mitochondrial reserve provides the energy required for the rapid induction of glycolysis upon T cell activation (van der Windt et al., 2013). We show that ATR kinase inhibition with ceralasertib during days 5, 6, and 7 post-infection suppresses immune memory at approximately seven weeks, demonstrating that ATR kinase activity within this window is critical for the differentiation of functional, long-lived memory cells. That a mere three-day window of ATR inhibition is sufficient to permanently reduce mitochondrial mass in CD8^+^ T memory cells, and irreversibly impair memory formation, indicates that, at least in this context, mitochondrial dysfunction cannot be repaired and is a key critical determinant of immune memory.

This study underscores the need to prioritize dosing schedule as a foundational element of clinical trial design with DNA damage response inhibitors. Prior work demonstrated that short-course ATR inhibition (days 1–3) combined with hypofractionated radiotherapy (days 1–2) effectively potentiates antitumor CD8⁺ T cell responses, whereas prolonged or improperly timed ATRi administration can impair the expansion of antigen-specific CD8⁺ T cells and suppress initial effector CD8⁺ T cell responses (25, 27). Here, we extend these findings by showing that ATRi administration on days 5–7 after LCMV Armstrong infection permanently impairs the formation of CD8⁺ T cell memory. These results are impactful since multiple ATR inhibitors (including ART0380, ceralasertib, BAY1895344, M1774, RP-3500, and others) are currently being evaluated in over 30 ongoing clinical trials, many of which combine them with radiotherapy, chemotherapy, or immune checkpoint inhibitors, in oncology settings where immune-mediated tumor control is increasingly recognized as a determinant of durable response. Since even classic cytotoxic chemotherapy relies on the immune system for part of its efficacy, combinations of ATRi and cytotoxic chemotherapies will also suffer from improper ATRi scheduling (67, 68).

Mitochondrial biogenesis and oxidative phosphorylation frequently promote cancer progression and metastasis (69, 70), and our data suggest that ATRi may exacerbate aberrant mitochondrial fission, which may lead to reduced oxidative phosphorylation capacity and increased apoptosis in vulnerable cancer cells. Disseminated cancer cells in melanoma, breast, and renal cancers have increased expression of the mitochondrial biogenesis transcription factor PGC-1α which promotes mitochondrial mass (71-73). BRAF and KRAS driven tumors have increased Erk-dependent DRP1 serine-616 phosphorylation and mitochondrial fission (74-77). Furthermore, mitochondrial fission is associated with enhanced apoptosis (78-80). ATRi may exacerbate aberrant mitochondrial fission and thereby reduce oxidative phosphorylation and increase apoptosis in some cancers.

In summary, the effects of ATR kinase inhibition on both adaptive immunity and mitochondrial function were unanticipated by decades of genetic studies and only became evident with highly selective, clinical-grade pharmacologic inhibitors. These findings highlight that the therapeutic impact of ATRi is highly schedule- and context-dependent. Poorly optimized timing can impair CD8⁺ T cell memory and antitumor immune surveillance, thereby undermining efficacy and contributing to suboptimal or failed trials, particularly when combined with radiotherapy, immune checkpoint blockade, or vaccines. Conversely, carefully timed ATR inhibition may be harnessed as a selective vulnerability against cancers that rely on enhanced mitochondrial function for survival, without compromising immune responses. Our data emphasize the urgent need to prioritize rational scheduling in the design of clinical trials evaluating ATRi.

## MATERIALS AND METHODS

### Cell lines and tissue culture

Splenocytes and CD8^+^ T cells cultured *ex vivo* were grown in complete T cell media comprised of RPMI-1640 (Corning) supplemented with 10% fetal bovine serum (FBS, R&D Systems), 100 U/mL penicillin and 100 mg/mL streptomycin (Lonza), 1x MEM NEAA, 1 mM sodium pyruvate, 5 mM HEPES, and 45 μM β-mercaptoethanol (all Gibco). HeLa cells, purchased from ATCC, and mouse embryonic fibroblasts, from cryopreserved stocks previously generated as described in (81), were grown in glucose-free RPMI-1640 (Gibco) supplemented with 5.5 mM glucose, 10% FBS, 100 U/mL penicillin and 100 mg/mL streptomycin, 0.1 mM NEAA, 1 mM sodium pyruvate, and10 mM HEPES. B16-F10 parental cells (B16), purchased from ATCC, and B16-F10 cells expressing the LCMV Armstrong glycoprotein (B16-GP), provided by Haydn T. Kissick, PhD (Emory University), were grown in DMEM containing 10% FBS and 100U/mL penicillin plus 100mg/mL streptomycin. Glycoprotein expression in B16-GP cells was verified by flow cytometry analysis of ZsGreen1 fluorescence intensity. All cell treatments *ex vivo* and *in vitro* were performed with 5 µM ATRi (ceralasertib, AstraZeneca or AdooQ Bioscience), 5 µM CDK1i (Ro-3306, Selleck Chem), and/or Vehicle (0.05% DMSO), for the indicated durations.

### CD8^+^ T cell activation and expansion

Female C57BL/6 mice and pmel-1 (B6.Cg-Thy1a/Cy Tg(TcraTcrb)8Rest/J) TCR transgenic mice were purchased from The Jackson Laboratory and were 6-8 weeks old at the time of spleen harvest for *ex vivo* culture experiments. For activation of CD8^+^ T cells from C57BL/6 mice, CD8^+^ T cells were isolated from splenocyte suspensions via negative selection using the EasySep Mouse CD8^+^ T Cell Isolation Kit (Stem Cell Technologies) according to the manufacturer’s instructions. Isolated CD8^+^ T cells were activated for 24 h in wells/dishes pre-coated overnight at 37°C with 10 µg/mL anti-CD3Ɛ (clone 145-2C11, BioLegend) and in complete T cell media containing 2 µg/mL anti-CD28 (clone 37.51, BD Pharmingen) and 50 U/mL IL-2 (Peprotech). For activation of CD8^+^ T cells from pmel-1 transgenic TCR mice, splenocytes were cultured for 24 h in complete T cell media containing 1 µM human gp100 peptide (aa 25-33, Anaspec) and 50 U/mL IL-2. Post activation, CD8^+^ T cells or splenocytes were collected, pelleted, resuspended in new complete T cell media containing 50 U/mL IL-2, and expanded for the indicated times. While the duration of was expansion varied across experiments, activation of naïve CD8^+^ T cells was always performed for 24 h using the same antibody or peptide concentrations and the same IL-2 concentration.

### pDRP1 immunoblotting and flow cytometry

*Ex vivo* activated CD8^+^ T cells isolated from C57BL/6 mouse spleens or splenocytes from Pmel-1 mice were treated for in 96-well round-bottom plates for 30 min with ATRi, CDK1i, combination (A+C), or Vehicle in 200 µL complete T cell media. Cells were then washed in 1x PBS, resuspended in 100 µL 1x PBS, and immediately fixed by adding 100 µL pre-warmed (37°C) Phosflow Fix Buffer I (BD Biosciences) and incubating for 10 min at 37°C. Cells were washed in 1x PBS, permeabilized for 30 min on ice in 200 µL pre-chilled (-20°C) Phosflow Perm Buffer III (BD Biosciences), and stored at -20°C for 1-3 days.

Cells were pelleted to remove Perm Buffer III, washed twice with flow cytometry staining (FCS) buffer (eBioscience), and blocked for 10 min at 4°C with 5% normal mouse serum (Invitrogen) in 75 µL FSC buffer. Cells were stained for 1 h 15 min on ice with Phospho-DRP1 (Ser616) rabbit monoclonal antibody (clone D9A1, 1:800, Cell Signaling Technologies) in 100 µL FSC buffer, then stained for 1 h at room temperature with AlexaFluor 647-conjugated goat anti-rabbit IgG (H+L) secondary antibody (1:3000, Invitrogen) in 150 µL FSC buffer. For splenocytes from Pmel-1 mice, anti-CD8a Spark YG 593 (BioLegend) and anti-TCRβ AlexaFluor (BioLegend) were added (both 1:300) during this secondary antibody staining step. For each treatment condition, samples stained with secondary antibody only were included to determine background/non-specific fluorescence. Cells were then stained for 30 min at room temperature with FxCycle Violet Ready Flow Reagent (2 drops per sample, Invitrogen) in 200 µL FSC buffer immediately prior to flow cytometry. Washes between staining steps were performed in FSC buffer.

Data were acquired using a 4-laser CytoFLEX cytometer and CytExpert software (both Beckman Coulter) with ≥5 x 10^4^ CD8^+^ T cells in S phase (based on the DNA content histogram) acquired per sample. For splenocytes from Pmel-1 mice, single stained splenocytes (and matching unstained cells) were used for single-color compensation controls. Data analyses were performed using FlowJo v10 software. Cell cycle gating was determined using the Watson Pragmatic algorithm univariate model in FlowJo v10. The Median fluorescence intensities (MFI) used for the reported normalized MFI were corrected for background/non-specific fluorescent signal by subtracting the MFI of the secondary antibody-only control within each given treatment condition prior to normalization to the mean of Vehicle controls. The gating strategy is shown in Supplemental Fig S6.

### Microscopy

Exponentially dividing mouse embryonic fibroblasts and HeLa cells were treated with Vehicle or ATRi for 6 h, inhibitor was washed out, and cells were allowed to recover for 18 h. During the final 2 h of the recovery period, mitochondria were stained for 2 h at 37°C with 2 ng/mL MitoView Fix 640 (Biotium) in complete media. Cells were then rinsed with 1x PBS, fixed for 30 min at room temperature in 2% paraformaldehyde in 1x PBS, and stained for 1 min at room temperature with 10 µg/ml Hoescht in 1x PBS. Samples were imaged in 1x PBS using a Nikon AX NSPARC system with NSPARC super-resolution mode and 60X 1.4NA Plan Apo objective. Channels were imaged sequentially, and a 2.5 µm Z-stack with a 0.19 µm step size was collected for each image. An Extended Depth of Focus image was created as a final image.

### Mitotracker Green flow cytometry in CD8^+^ T cells activated *ex vivo*

*Ex* vivo activated CD8^+^ T cells isolated from C57BL/6 mouse spleens were treated for 6 h with ATRi or Vehicle and then seeded in 96-well round-bottom plates. Cells were stained for 20 min at 37°C with MitoTracker Green FM for flow cytometry (1:1000, Invitrogen) in 75 µL 1x PBS. Cells were stained with for 10 min at 4°C with eFluor 780 viability dye (1:2000) in 50 µL 1x PBS. Washes between staining steps were performed in 2% FBS in 1x PBS. Data were acquired from live, unfixed cells using a 4-laser CytoFLEX cytometer and CytExpert software (both Beckman Coulter) with ≥4 x 10^4^ live, CD8^+^ T cells acquired per sample. Data analyses were performed using FlowJo v10 software.

### Metabolic flux analysis

Activated CD8^+^ T cells were treated with Vehicle or ATRi for 6 h either prior to an 18 h recovery period or following an 18 h expansion period, as indicated. At 48 h hours, cells were collected, pelleted, and resuspended in Seahorse XF assay media (Agilent) supplemented with 10 mM glucose, 2 mM glutamine, and 1 mM sodium pyruvate. Cells were counted via hemocytometer and 10^5^ live cells (trypan blue negative) were seeded in eight replicate wells (unless otherwise stated in the legend) for metabolic flux analysis using the in XFe96/XF Pro PDL FluxPak Mini, Seahorse XFe96 Analyzer, and Seahorse Wave software (all Agilent). Oxygen consumption rate (OCR, pmol/min) and extracellular acidification rate (ECAR, mpH/min) were measured every 6.9 minutes for 98 minutes. Injections of 2 µM oligomycin (Oligo, Caymen Chemical), 2 µM FCCP (Caymen Chemical), 10 mM 2-deoxyglucose (2-DG, Sigma), and 0.5 µM each of rotenone and antimycin A (Rot/AA, both Sigma) were delivered sequentially at the indicated intervals. Spare respiratory capacity was calculated as maximal OCR - basal OCR (OCR_max_-OCR_basal_). Reported OCR, ECAR, and spare respiratory values represent the means (± SEM) of background-corrected technical replicate values, as calculated and provided by the Agilent Seahorse Analytics XF Software.

### Mice and treatments

Female C57BL/6 mice were purchased from The Jackson Laboratory and were 6-8 weeks old at the time of infection with 1.5 x 10^6^ PFU LCMV Armstrong (in 150 µL), administered intraperitoneally on day 0. Uninfected control mice of the same age and housed in the same room were included with every LCMV experiment. For all *in vivo* treatments, mice received 75 mg/kg ATRi (ceralasertib) or Vehicle (10% DMSO, 40% Propylene Glycol, 50% dH_2_O) via oral gavage on the days indicated. For antigen rechallenge experiments, 2.5 x 10^5^ B16-GP (or parental B16) cells in 50 µL 1x PBS were injected subcutaneously into both flanks of mice 6 weeks after LCMV-Armstrong infection. Tumors were measured twice weekly with calipers, and volumes were calculated as volume = (length x width)^2^/2. The survival endpoint was reached when tumor length exceeded 15 mm, when tumors ulcerated, or when pressure necrosis preceding ulceration was evident.

### Tissue processing

Spleens were harvested at the indicated time points post LCMV infection or at 6-8 weeks of age (for *ex vivo* culture experiments). Spleens were mechanically dissociated with a 3 mL syringe plunger or with frosted glass slides and were filtered through 70 μm cell strainers (Corning). Erythrocytes were lysed in 1 mL of 150 mM NH_4_Cl, 10 mM NaHCO^3^, 0.1 mM EDTA (pH 8.0) for 30-40 seconds. Lysis was quenched by addition of ≥ 5 mL serum-free RPMI-1640 media. Cells were pelleted, resuspended in complete T cell media, and filtered again through 70 μm cell strainers. Splenocyte suspensions were counted using a Scepter 3.0 handheld counter (Millipore).

### Immune profiling by multiparameter spectral flow cytometry

Splenocytes were seeded at 2 x 10^6^ cells in 96-well round-bottom plates for staining. Fc receptors were blocked for 10 minutes at 4°C with of 0.5 μg anti-CD16/32 antibody (TruStain FcX Plus, BioLegend) in 100 µL FSC buffer (Invitrogen eBioscience Flow Cytometry Staining Buffer). LCMV GP33-specific CD8^+^ T cells were labeled for 30 minutes at 4°C with APC-conjugated GP33 (LCMV gp 33-41 KAVYNFATC or KAVYNFATM) H2-Db tetramer (1:100, NIH Tetramer Facility) at in 50 µL FSC buffer. Cells were then stained for 15-30 minutes at 4°C with antibodies against surface antigens in 50 µL FSC buffer with Brilliant Stain Buffer Plus (1:5, BD Biosciences) added to prevent polymer dye-dye interactions in antibody cocktails containing multiple Brilliant Violet dye conjugates. After surface staining, dead/dying cells were stained for 10 minutes at 4°C with eFluor 780 viability dye (1:2000; Invitrogen) in 50 µL 1x PBS. Samples were then fixed and permeabilized for 15 minutes at room temperature in 75 µL eBioscience Fixation/Permeabilization reagent (Invitrogen) and were stained for 45 minutes at room temperature with antibodies against nuclear proteins (Ki67, TCF1, T-bet) in 50 µL eBioscience 1x Permeabilization Buffer (Invitrogen). Washes between staining steps were performed pre-fixation in FSC buffer and post-fixation in 1x Permeabilization Buffer.

For profiling of GP33-specific memory CD8^+^ T cell functionality (IFN-γ, TNF-α production) and proliferative capacity (Ki67), splenocytes in 96-well round bottom plates from previously LCMV-infected (and uninfected control) mice were stimulated in 200 µL complete T cell media with 100 ng/mL GP33 peptide for 4 h or with 100 ng/mL GP33 and 25 U/mL for 22 h, with 5x eBioscience Protein Transport Inhibitor Cocktail (Invitrogen) in 50 µL T cell media added to the existing media (for a final 1x concentration) for the final 4 h of stimulation. Fc receptor blocking, viability dye staining, and fixation/permeabilization were performed as described above, while surface and intracellular/nuclear antibody staining was performed concurrently post fixation/permeabilization. For assessing proliferative capacity (expansion) after stimulation, cells were stimulated in complete T cell media with 100 ng/mL GP33 peptide and 50 U/mL IL-2 for 18 h and were then expanded in complete T cell media with 50 U/mL IL-2 for an additional 72 h. Cells were then seeded into 96-well round-bottom plates for Fc receptor blocking, surface antibody staining, viability dye staining, and fixation/permeabilization, performed as described above.

For profiling of mitochondrial mass in GP33-specific memory CD8^+^ T cells, splenocytes (2 x 10^6^ cells) from treated mice were harvested at 6 weeks post LCMV infection and stained in 96-well round-bottom plates for 15 min at 37°C with MitoTracker Green FM for flow cytometry (1:1000, Invitrogen) at 100 µL serum-free T cell media (RPMI-1640 supplemented with 100 U/mL penicillin and 100 mg/mL streptomycin, 1x MEM NEAA, 1 mM sodium pyruvate, and 5 mM HEPES, and without FBS and β-mercaptoethanol). Next, Fc receptors were blocked for 10 minutes at 4°C with 0.5 μg anti-CD16/32 antibody (TruStain FcX Plus, BioLegend) in 100 µL FSC buffer (Invitrogen eBioscience Flow Cytometry Staining Buffer), LCMV GP33-specific CD8^+^ T cells were labeled for 30 minutes at 4°C with PE-conjugated GP33 (LCMV gp 33-41 KAVYNFATM) H2-Db tetramer (1:100, NIH Tetramer Facility) in 50 µL FSC buffer, and cells were stained for 15 minutes at 4°C with antibodies against surface antigens (listed in Supplemental Table S1) in 50 µL FSC buffer with Brilliant Stain Buffer Plus, and dead/dying cells were stained for 10 minutes at 4°C with eFluor 780 viability dye (1:3000, Invitrogen) in 100 µL 1x PBS. Washes between staining steps were performed in 2% FBS in 1x PBS. Data were collected from live, unfixed cells with ≥5000 live, GP33^+^ CD8^+^ T cells acquired per sample.

Data for all spectral cytometry experiments were acquired using a Cytek Biosciences Aurora spectral flow cytometer, with spectral unmixing and data acquisition performed in the Cytek Biosciences SpectroFlo software. Single stained splenocytes (and matching unstained cells) or single stained UltraComp eBeads (Invitrogen) were used for single-color compensation controls. Uninfected controls, unstimulated controls, and/or fluorescence minus one (FMO) controls were used, where appropriate, to empirically determine gating. Analyses of unmixed data were performed using FlowJo v10 software. Antibody information (target, clone, dilution, manufacturer) for the staining panels is included in Supplemental Table 1. Gating strategies are shown in Supplemental Figures S7-S9.

### RNA sequencing analyses

Bulk RNA sequencing was performed on sorted GP33-specific (GP33) CD8^+^ T cells at 6 weeks post-infection. Spleens were harvested and processed as described above, and CD8^+^ T cells were isolated via negative selection using the EasySep Mouse CD8^+^ T Cell Isolation Kit (Stem Cell Technologies) according to the manufacturer’s instructions. CD8^+^ T cell suspensions were counted using a Scepter 3.0 handheld counter (Millipore) and seeded at 10^6^ cells in 96-well round-bottom plates for staining. Fc receptors were blocked and cells were stained with APC-conjugated GP33 loaded-tetramer, anti-CD8a, and eFluor 780 viability dye, as described above. Live cells (500-1,000 cells/well) were sorted with a BigFoot Spectral Cell Sorter (ThermoFisher) into a 96-well plate containing cell lysis buffer. The plate was stored at -80°C until the University of Pittsburgh Health Sciences Sequencing Core performed RNA extraction, cDNA library preparation, and sequencing.

RNA sequencing data (fastq) underwent quality control assessment using FastQC (82) to evaluate sequence quality metrics. Following quality control, sequencing reads were aligned to the mouse reference genome (GRCm39/mm39, Ensembl release 104) using the STAR algorithm (v2.7.1) (83) within the RSEM workflow (v1.3.3) (84). This integrated approach facilitated the generation of both raw read counts and Transcripts Per Million (TPM) expression values for each transcript. TPM values were normalized using the formula log₂(TPM + 1) to stabilize variance and approximate a normal distribution for downstream statistical analyses. To identify differentially expressed genes (DEGs) between the vehicle- and ATRi-treated groups, the mean log₂(TPM+1) expression values were calculated for each gene in each group. Gene level log₂ fold-changes (log₂FC) were computed as the difference in mean log₂(TPM+1) expression between the groups (ATRi-treated minus vehicle-treated). An unadjusted two-sample t-test was then performed on the log₂(TPM+1) values to assess differences in expression between groups. Genes with absolute fold changes in expression ≥1.5 and p-values ≤0.01 were considered differentially expressed for exploratory analysis; no multiple testing correction was applied at this stage.

### Statistics

For survival analyses, significance was determined by log-rank test with Holm-Šidák adjustment for multiple comparisons. For longitudinal tumor growth, significance was determined by mixed effects model comparing LCMV-infected treatment groups, with Holm-Šidák adjustment for multiple comparisons when multiple comparisons were made. For both survival analyses and longitudinal tumor growth, all comparisons made are denoted by brackets in the figures. For splenocyte immune profiling, MitoTracker Green experiments, and pDRP1 experiments, statisitcal significance was determined one-way ANOVA with Tukey’s multiple comparisons test when comparing multiple groups, and by unpaired, two-tailed Welch’s t-test when comparing only two groups (Vehicle versus ATRi). For metabolic flux experiments (Searhose XF Analayzer), statisitcal significance was determined one-way ANOVA with Šidák ‘s multiple comparisons test when comparing multiple groups, and by unpaired, two-tailed Welch’s t-test when comparing only two groups (Vehicle versus ATRi). All statistically significant were shown unless otherwise noted in the legend. A 95% confidence interval and significance of p<0.05 were used for all statistical tests, except that Holm-Šidák adjustments for multiple comparisons do not compute confidence intervals. All statistical analyses were performed in GraphPad Prism 10.

