## SUPPLEMENTAL DATA for "ATR kinase inhibitors induce mitochondrial fission in CD8^+^ T cells and impair immune memory *in vivo*"

**Supplemental Methods**

**pDRP1 flow cytometry in HEK 293T cells**

HEK 293T cells were purchased from ATCC and cultured in DMEM (Corning) supplemented with 10% fetal bovine serum (FBS, R&D Systems) and 100 U/mL penicillin and 100 mg/mL streptomycin (Lonza). Exponentially-dividing HEK 293T cells were treated for 30 min with 5 µM ATRi (ceralasertib), 5 µM CDK1i (Ro-3306), combination (A+C), or Vehicle (Veh, 0.05% DMSO). Cells were washed with 1x PBS, dissociated with TrypLE Express (phenol red-free, Gibco), collected into complete media, pelleted, resuspended in 500 µL 1x PBS, and immediately fixed by adding 500 µL of pre-warmed (37°C) Phosflow Fix Buffer I (BD Biosciences) and incubating for 10 min at 37°C. Cells were washed in 1x PBS, permeabilized for 30 min on ice in 1 mL µL pre-chilled (-20°C) Phosflow Perm Buffer III (BD Biosciences), and stored at -20°C for 1-3 days. Cells were pelleted to remove Perm Buffer III, washed twice with flow cytometry staining (FCS) buffer (eBioscience), and blocked for 10 min at 4°C with 5% normal human serum (Invitrogen) in 100 µL (1st experiment) or 200 µL FSC buffer (2nd experiment). Cells were stained for 30 min (1st experiment) or 45 min (2nd experiment) on ice with Phospho-DRP1 (Ser616) rabbit monoclonal antibody (clone D9A1, 1:800, Cell Signaling Technologies) in 50 µL FSC buffer, then stained for 40 min at room temperature with AlexaFluor 647-conjugated goat anti-rabbit IgG (H+L) secondary antibody (Invitrogen, 1st experiment 1:1000, 2nd experiment 1:2000) in 50 µL FSC buffer. For each treatment condition, samples stained with secondary antibody only were included to determine background/non-specific fluorescence. Cells were then stained for 15 min at room temperature with 200 µL FxCycle PI/RNase Staining Solution (Invitrogen, 1st experiment) or 20 min at room temperature with FxCycle Violet Ready Flow Reagent

(1 drop per sample, Invitrogen) in 200 µL FSC buffer immediately prior to flow cytometry. Washes between staining steps were performed in FSC buffer. Data were acquired using a 4-laser CytoFLEX cytometer and CytExpert software (both Beckman Coulter) with >$10^4$ cells analyzed per sample. Data analyses were performed using FlowJo v10 software. The Median fluorescence intensities (MFI) used for the reported normalized MFI were corrected for background/non-specific fluorescent signal by subtracting the MFI of the secondary antibody-only control within each given treatment condition prior to normalization to the mean of Vehicle controls.

**CD8$^+$ T cell activation and expansion**

CD8$^+$ T cells were isolated from splenocyte suspensions from 6-8-week-old female C57BL/6 mice (Jackson Laboratory) via negative selection using the EasySep Mouse CD8$^+$ T Cell Isolation Kit (Stem Cell Technologies) according to the manufacturer's instructions. Isolated CD8$^+$ T cells were activated for 24 h in wells/dishes pre-coated overnight at 37°C with 10 µg/mL anti-CD3Ɛ (clone 145-2C11, BioLegend) and in complete T cell media (RPMI-1640 supplemented with 10% fetal bovine serum, 100 U/mL penicillin and 100 mg/mL streptomycin, 1x MEM NEAA, 1 mM sodium pyruvate, 5 mM HEPES, 45 µM β-mercaptoethanol) containing 2 µg/mL anti-CD28 (clone 37.51, BD Pharmingen) and 50 U/mL IL-2 (Peprotech). Post activation, CD8$^+$ T cells or splenocytes were collected, pelleted, resuspended in new complete T cell media containing 50 U/mL IL-2, and expanded for the indicated times.

**Mitotracker Green flow cytometry in CD8$^+$ T cells activated *ex vivo***

*Ex vivo* activated CD8$^+$ T cells isolated from C57BL/6 mouse spleens were treated for 6 h with ATRi or Vehicle after either 24 h or 48 h expansion in IL-2 and were then either immediately seeded in 96-

well round-bottom plates or were allowed to recover to for 18 h absent ATRi prior to seeding, respectively. Cells were stained for 15-20 min at 37°C with MitoTracker Green FM for flow cytometry (1:1000, Invitrogen) in 100 µL 1x PBS or 100 µL serum-free T cell media (RPMI-1640 supplemented with 100 U/mL penicillin and 100 mg/mL streptomycin, 1x MEM NEAA, 1 mM sodium pyruvate, and 5 mM HEPES, and without FBS and β-mercaptoethanol). Cells were stained with for 10 min at 4°C with eFluor 780 viability dye (1:2000-1:3000) in 50 µL 1x PBS. Washes between staining steps were performed in 2% FBS in 1x PBS. Data were acquired from live, unfixed cells using a 4-laser CytoFLEX cytometer and CytExpert software (both Beckman Coulter) with >2.5 x $10^4$ live, $CD8^+$ T cells acquired per sample. Data analyses were performed using FlowJo v10 software. Median fluorescence intensities (MFI) were normalized to the mean of Vehicle controls.

**Metabolic flux analysis**

Activated $CD8^+$ T cells were treated with Vehicle or ATRi ± 6 µM thymidine (dT) for 6 h following an 18 h expansion period. At 48 h hours, cells were collected, pelleted, and resuspended in Seahorse XF assay media (Agilent) supplemented with 10 mM glucose, 2 mM glutamine, and 1 mM sodium pyruvate. Cells were counted via hemocytometer and $10^5$ live cells (trypan blue negative) were seeded in eight replicate wells for metabolic flux analysis using the in XFe96/XF Pro PDL FluxPak Mini, Seahorse XFe96 Analyzer, and Seahorse Wave software (all Agilent). Oxygen consumption rate (OCR, pmol/min) and extracellular acidification rate (ECAR, mpH/min) were measured every 6.9 minutes for 98 minutes. Injections of 2 µM oligomycin (Oligo, Caymen Chemical), 2 µM FCCP (Caymen Chemical), 10 mM 2-deoxyglucose (2-DG, Sigma), and 0.5 µM each of rotenone and antimycin A (Rot/AA, both Sigma) were delivered sequentially at the indicated intervals. Spare respiratory capacity was calculated as maximal OCR - basal OCR ($OCR_{max}$-$OCR_{basal}$). Reported

OCR, ECAR, and spare respiratory values represent the means (± SEM) of background-corrected technical replicate values, as calculated and provided by the Agilent Seahorse Analytics XF Software.

**Supplemental Table S1.** Antibodies for Flow Cytometry

| Antibody/Tetramer | Clone | Dilution | Manufacturer | Catalog # |
|---|---|---|---|---|
| anti-phospho-DRP1 (Ser616) | D9A1 | 1:800 | Cell Signaling Technologies | 4494S |
| TruStain FcX PLUS anti-mouse CD16/32 | S17011E | 1:100 | BioLegend | 156604 |
| APC GP33 (LCMV gp 33-41 KAVYNFATM) H2-Db tetramer | | 1:100 | NIH Tetramer Facility | |
| APC GP33 (LCMV gp 33-41 KAVYNFATC) H2-Db tetramer | | 1:100 | NIH Tetramer Facility | |
| PE GP33 (LCMV gp 33-41 KAVYNFATM) H2-Db tetramer | | 1:100 | NIH Tetramer Facility | |
| AF488 anti-CD8a | KT15 | 1:40 | BioRad | MCA609A488 |
| AF488 anti-TCR-β | H57-597 | 1:300 | BioLegend | 109216 |
| AF647 anti-CD8a | KT15 | 1:40 | BioRad | MCA609A647 |
| AF647 goat anti-rabbit IgG (H+L) | polyclonal | 1:2000-1:3000 | Invitrogen | A21244 |
| AF647 anti-IFN-γ | XMG1.2 | 1:500 | BioLegend | 505816 |
| BV421 anti-CD4 | RM4-5 | 1:600 | BioLegend | 100563 |
| BV421 anti-TNF-α | MP6-XT22 | 1:250 | BioLegend | 506328 |
| BV510 anti-CD4 | GK1.5 | 1:300 | BioLegend | 100449 |
| BV510 anti-CD8a | 53-6.7 | 1:200 | BioLegend | 100752 |
| BV605 anti-CD62L | MEL-14 | 1:300 | BioLegend | 104438 |
| BV650 anti-Ki67 | B56 | 1:100 | BD Biosciences | 563757 |
| BV711 anti-CD25 | PC61 | 1:300 | BioLegend | 102049 |
| BV785 anti-TIM3 | RMT3-23 | 1:100 | BioLegend | 119725 |
| BUV395 anti-CD45 | 30-F11 | 1:250-1:300 | BD Biosciences | 564279 |
| BUV563 anti-TCR-β | H57-597 | 1:300 | BD Biosciences | 748406 |
| BUV563 anti Thy1.2 | 30-H12 | 1:300 | BD Biosciences | 741214 |
| BUV737 anti-CD44 | IM7 | 1:500 | BD Biosciences | 612799 |
| Pacific Blue anti-TCF1/TCF7 | C63D9 | 1:100 | Cell Signaling Technology | 9066S |
| PerCP-Cy5.5 anti-T-bet | 4B10 | 1:100 | BioLegend | 644806 |
| PerCP-eFluor 710 anti-KLRG1 | 2F1 | 1:300 | eBioscience | 46-5893-80 |
| PE anti-CD69 | H1.2F3 | 1:300 | eBioscience | 12-0691-82 |
| PE-Cy7 anti-CD127 | A7R34 | 1:300 | Tonbo Biosciences | 60-1271-U100 |
| PE-Dazzle 594 anti-PD-1 | RMP1-30 | 1:400 | BioLegend | 109116 |
| Spark YG 593 anti-CD8a | 53-6.7 | 1:300 | BioLegend | 285424 |

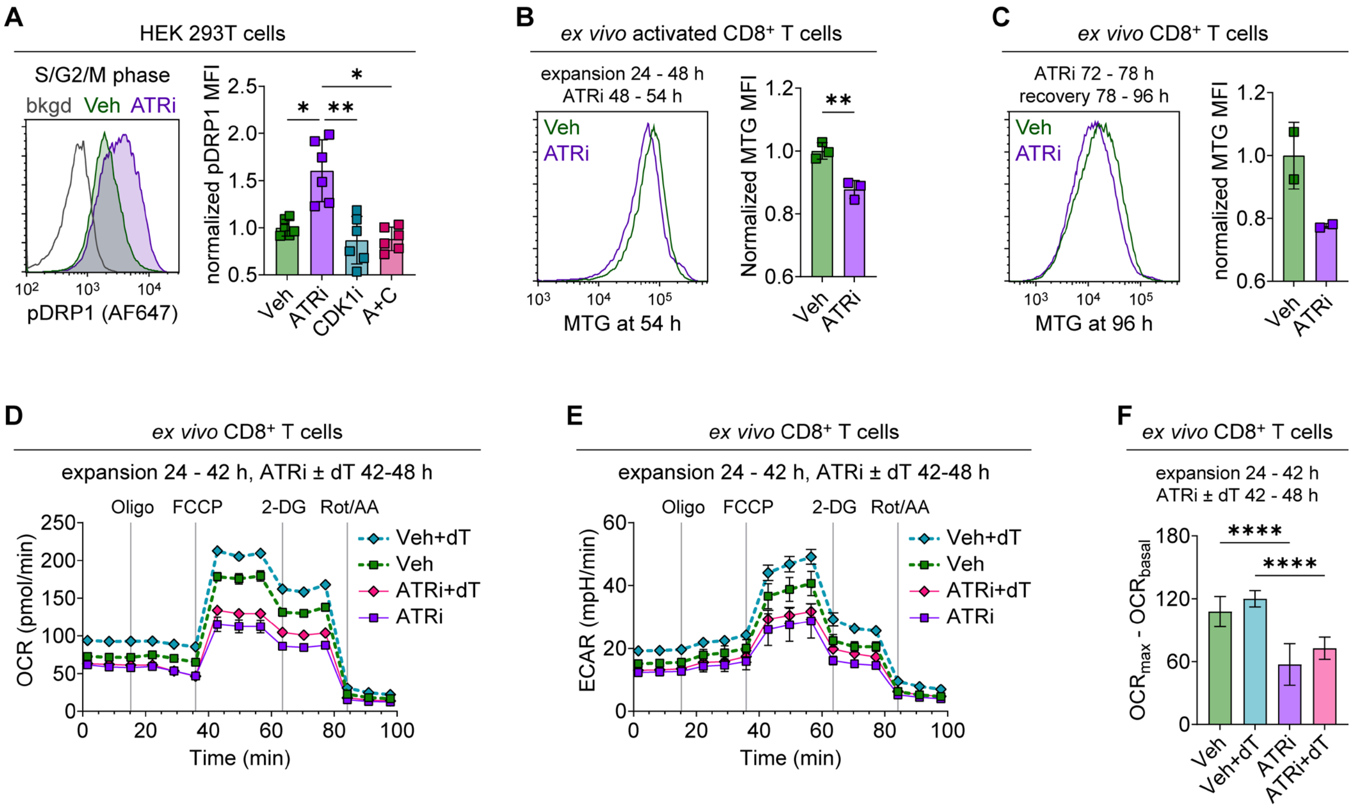


**Supplemental Figure S1. ATRi dysregulates DRP1 activity, reduces mitochondrial mass, and impairs bioenergetic capacity.**

**A.** HEK 293T cells were treated with Vehicle (Veh) or 5 µM ATRi ± 5 µM CDK1i for 30 min and then stained for flow cytometry analyses of DRP1 phosphorylation and DNA content. Shown are representative histograms of pDRP1 fluorescence intensity, and background (bkgd) fluorescence (secondary antibody only control), in non-G1 (S/G2/M) phase cells, as well as quantitation of background-corrected pDRP1 median fluorescence intensity (MFI), normalized to the mean of Veh controls, for non-G1 (S/G2/M) phase cells, as determined by DNA content. Data points with mean ± SD bars shown. Data combined from two independent experiments, each with three biological replicates. *p<0.05, **p<0.01 by one-way ANOVA with Tukey's multiple comparisons test. **B-F.** $CD8^+$ T cells were isolated (via negative selection) from wildtype C57BL/6 spleens and were activated *ex vivo* with plate-bound anti-CD3 (10 µg/mL), soluble anti-CD28 (2 µg/mL) antibodies, and 50 U/mL IL-2 for 24 h. **B.** Activated $CD8^+$ T cells were expanded for 24 h, treated with Veh or 5 µM ATRi for 6 h, and then stained with MitoTracker Green (MTG). Shown are representative histograms of MTG fluorescence intensity and quantitation of MTG MFI, normalized to the mean of Veh controls. Data from one experiment with three separately activated replicates (from two pooled spleens) per group. Mean ± SD bars shown. **$p<0.01$ by unpaired, two-tailed Welch's t-test. **C.** Activated $CD8^+$ T cells were expanded for 48 h and then treated with Veh or 5 μM ATRi for 6 h. Inhibitor was washed out and cells were allowed to recover for an additional 18 h prior to staining with MTG. Shown are representative histograms of MTG fluorescence intensity and quantitation of MTG MFI, normalized to the mean of Veh controls. Data from one experiment with two separately activated replicates (from three pooled spleens) per group. Individual data points represent the average of technical duplicates. Mean ± SD bars shown. Since $n < 3$, statistical analysis was not performed. **D-F.** Activated $CD8^+$ T cells were expanded for 18 h and then treated with Veh or 5 μM ATRi ± 6 μM thymidine (dT) for 6 h prior to metabolic flux analyses using the Seahorse XF Analyzer. **D.** Oxygen consumption rate (OCR, pmol/min) over time **E.** Extracellular acidification rate (ECAR, mpH/min) over time. **F.** Quantitation of spare respiratory capacity, calculated as maximal OCR - basal OCR. Data from one experiment. Bars represent the mean of eight technical replicates, with SEM bars shown. ****$p<0.0001$ by one-way ANOVA with Šidák 's multiple comparisons

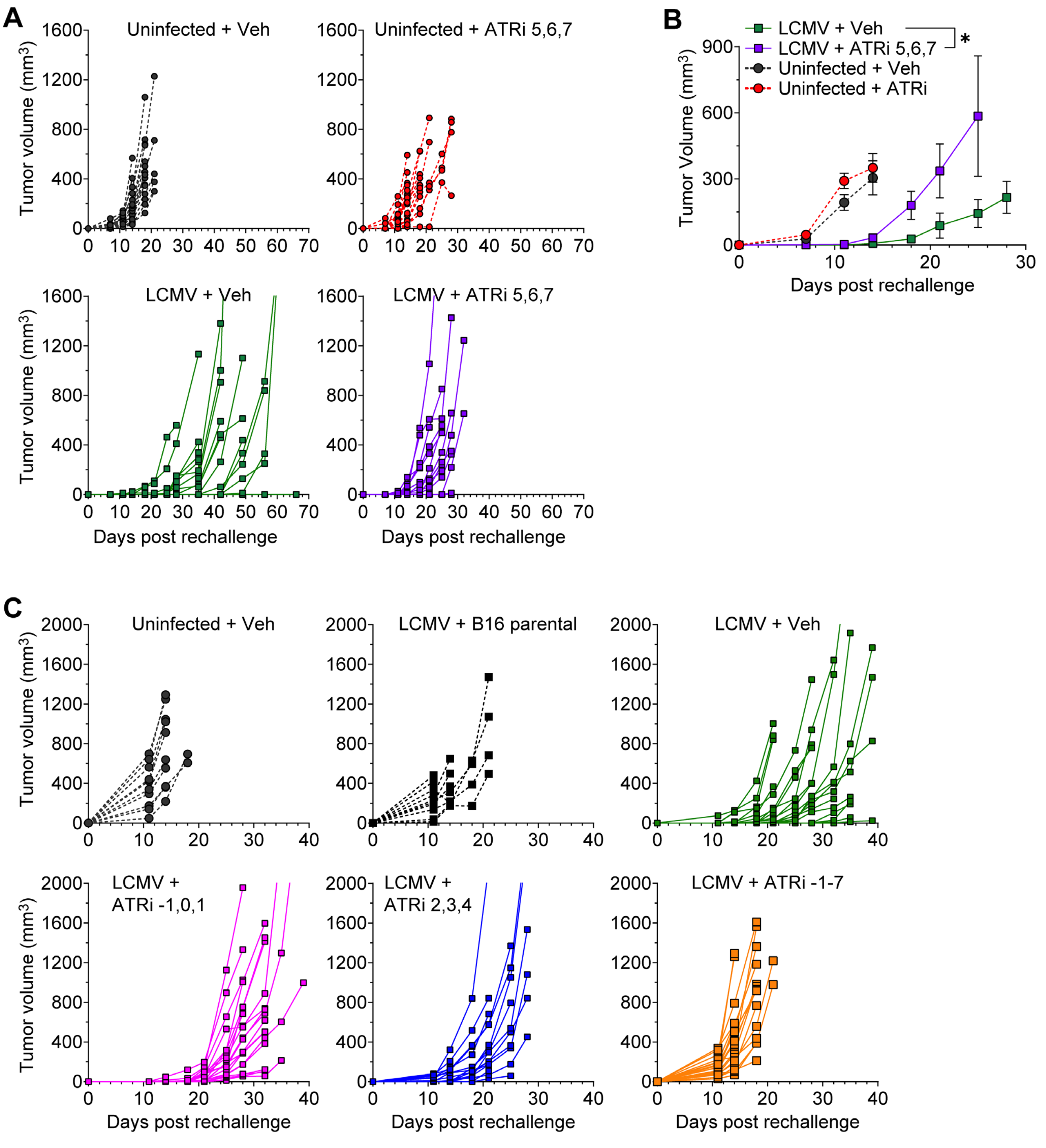


**Supplemental Figure S2. ATR signaling is essential to establish CD8⁺ T cell memory.**

**A.** Individual tumor growth curves for uninfected control groups and LCMV Armstrong-infected groups (from experiment shown in Figure 3B-C) treated daily with Vehicle (Veh) or 75 mg/kg ATRi on days 5, 6, and 7 post LCMV-infection. Data from one experiment with n = 7 mice (14 tumors) for

LCMV + ATRi 5,6,7 and n = 10 mice (20 tumors) for all other groups. **B.** Repeat of the rechallenge experiment in Figure 3B-C, but with tumors in one flank only. Shown are tumor growth curves with mean tumor volume ± SEM. Data from one experiment with n = 10 mice per group. *$p<0.05$ by mixed effects model comparing the LCMV-infected groups. **C.** Individual tumor growth curves for control groups and LCMV Armstrong-infected treatment groups (from experiment shown in Figure 3D-E) treated with Veh (days 2, 3, 4) or different schedules of 75 mg/kg ATRi, as indicated (days -1, 0, 1; days 2, 3, 4; or days -1 to 7). Controls include Veh-treated, uninfected mice injected with B16-GP tumors and Veh-treated, LCMV-infected mice injected with parental B16 (non-GP-expressing) tumors. Data from one experiment with n = 5 mice (10 tumors) for Uninfected + Vehicle and LCMV + B16 Parental; n = 8 mice (16 tumors) for LCMV + ATRi 2,3,4; and n = 10 mice (20 tumors) for all remaining groups.

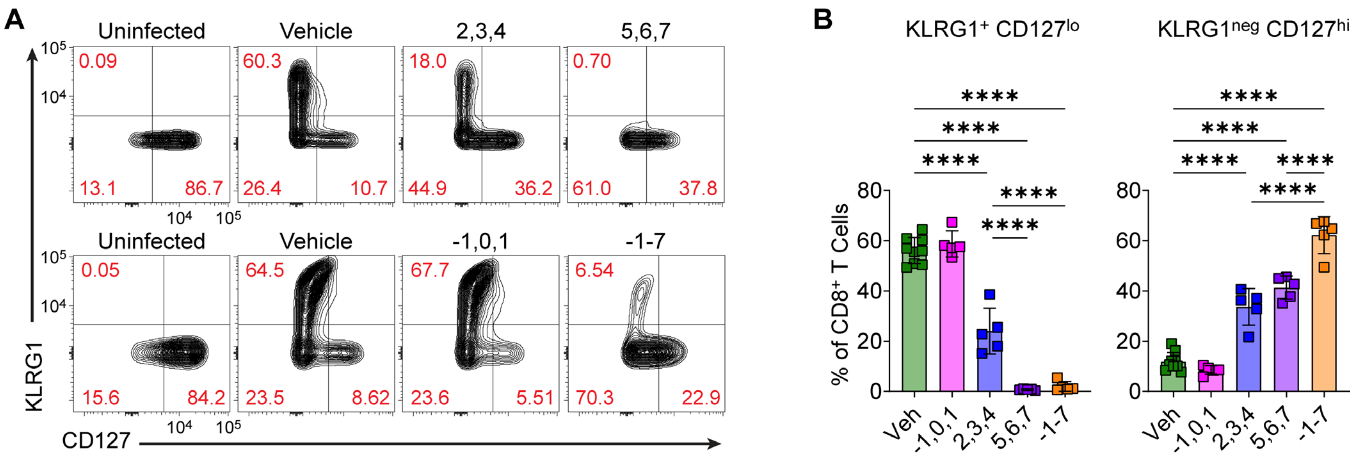


**Supplemental Figure S3. ATR signaling is essential for antigen-specific effector CD8$^+$ T cell clonal expansion.**

Mice were infected with LCMV Armstrong on day 0 and were treated daily with Vehicle (Veh, days 2, 3, 4 or days 5, 6, 7) or with different schedules of 75 mg/kg ATRi as indicated (days -1, 0, 1; days 2, 3, 4; or days -1 to 7). Splenocytes were immune profiled by spectral flow cytomety on day 8. **A.** Representative contour plots showing KLRG1 and CD127 expression on CD8$^+$ T cells from uninfected control, Veh-treated, and ATRi-treated mice (treatment days indicated), with the top row from one experiment and the bottom row from a second experiment. **B.** Quantification of KLRG1$^+$CD127$^{lo}$ and KLRG1$^{neg}$CD127$^{hi}$ CD8$^+$ T cells, as percentages of total CD8$^+$ T cells. Individual data points with mean ± SD bars shown. Data combined from two experiments, each with n = 5 mice per group per experiment, for a total n = 10 Veh and n = 5 for each ATRi treatment group. One uninfected control was included in each experiment. ****p<0.0001 by one-way ANOVA with Tukey's multiple comparisons test. Statistically significant comparisons not shown for -1,0,1 versus other ATRi groups (2,3,4 or 5,6,7 or -1 to 7), but are identical to Veh versus those ATRi groups. All other statistically significant comparisons are shown.

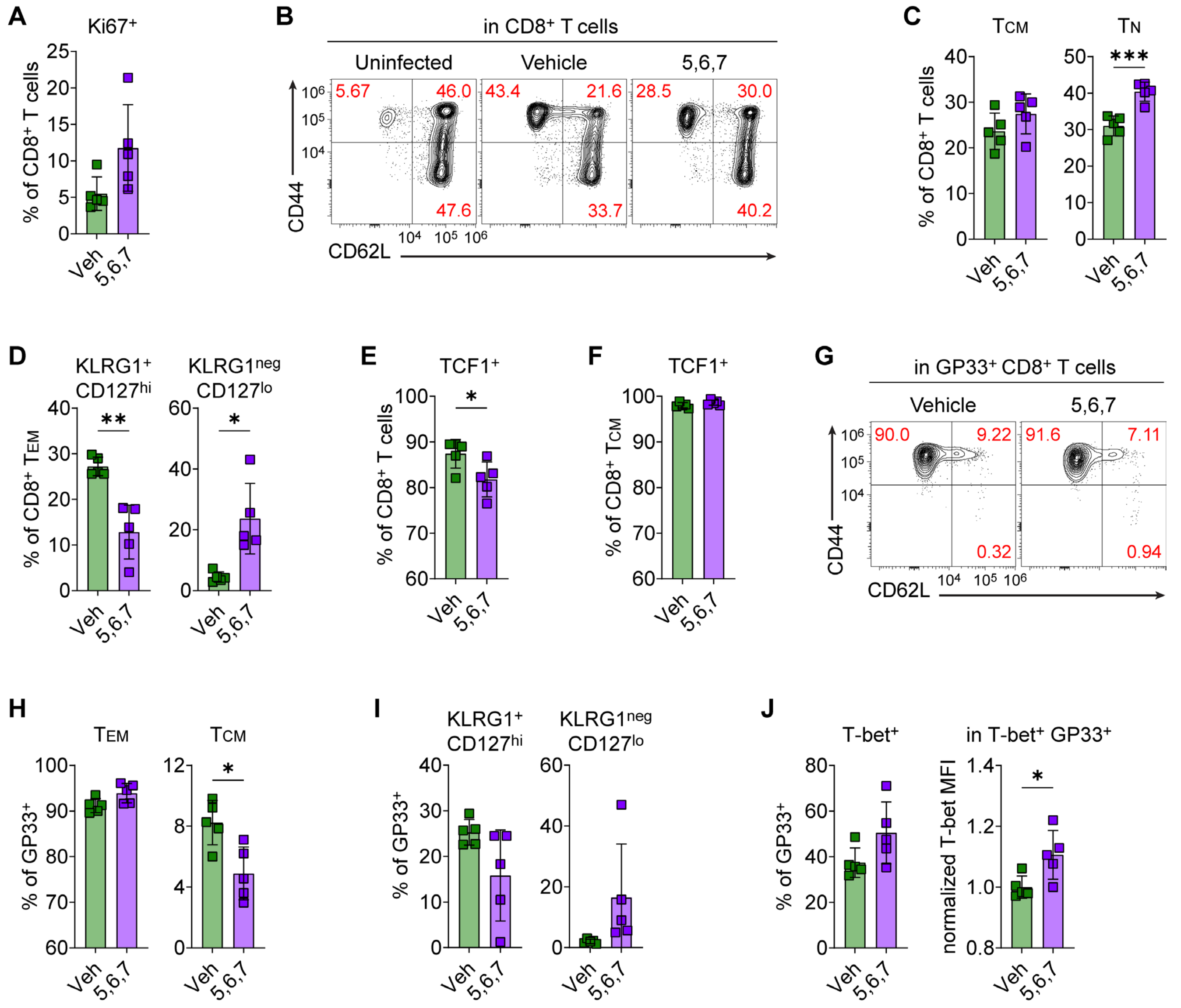


**Supplemental Figure S4. Cessation of ATRi triggers a delayed effector $CD8^+$ T cell response.** Mice were infected with LCMV Armstrong on day 0 and were treated daily with Vehicle (Veh) or 75 mg/kg ATRi (days 5, 6, 7). Splenocytes were immune profiled by spectral flow cytomety on day 18. **A.** Quantitation of the $Ki67^+$ $CD8^+$ T cells, as a percentage of total $CD8^+$ T cells. **B.** Representative contour plots showing CD44 and CD62L expression on $CD8^+$ T cells from uninfected control, Veh-treated, and ATRi-treated mice **C.** Quantitation of $CD8^+$ TCM ($CD44^+CD62L^+$) and $CD8^+$ TN ($CD44^{neg}CD62L^+$), as percentages of total $CD8^+$ T cells. **D.** Quantification of $KLRG1^+CD127^{hi}$ (ie. double positive) and $KLRG1^{neg}CD127^{lo}$ (ie. double negative) $CD8^+$ TEM, as percentages of total $CD8^+$ TEM. **E.** Quantification of $TCF1^+$ $CD8^+$ T cells, as a percentage of total $CD8^+$ T cells. **F.** Quantification of TCF1$^{+}$ CD8$^{+}$ T$_{CM}$, as a percentage of total CD8$^{+}$ T$_{CM}$. **G.** Representative contour plots showing CD44 and CD62L expression on GP33$^{+}$ CD8$^{+}$ T cells from Veh-treated and ATRi-treated mice. **H.** Quantitation of GP33$^{+}$ CD8$^{+}$ T$_{EM}$ (CD44$^{+}$CD62L$^{neg}$) and GP33$^{+}$ CD8$^{+}$ T$_{CM}$ (CD44$^{+}$CD62L$^{+}$), as percentages of total GP33$^{+}$ CD8$^{+}$ T cells. **I.** Quantification of KLRG1$^{+}$CD127$^{hi}$ (ie. double positive) and KLRG1$^{neg}$CD127$^{lo}$ (ie. double negative) GP33$^{+}$ CD8$^{+}$ T cells, as percentages of total GP33$^{+}$ CD8$^{+}$ T cells. **J.** Quantification of T-bet$^{+}$ GP33$^{+}$ CD8$^{+}$ T cells, as a percentage of total GP33$^{+}$ CD8$^{+}$ T cells, and quantitation of T-bet median fluorescence intensity (MFI), normalized to the mean of Veh-treated mice, in T-bet$^{+}$ GP33$^{+}$ CD8$^{+}$ T cells. **A-J.** Data from one experiment with n = 5 mice per group and including one uninfected control. **A, C-F, H-J.** Individual data points with mean ± SD bars shown. *p<0.05, **p<0.01, ***p<0.001 by unpaired, two-tailed Welch's t-test.

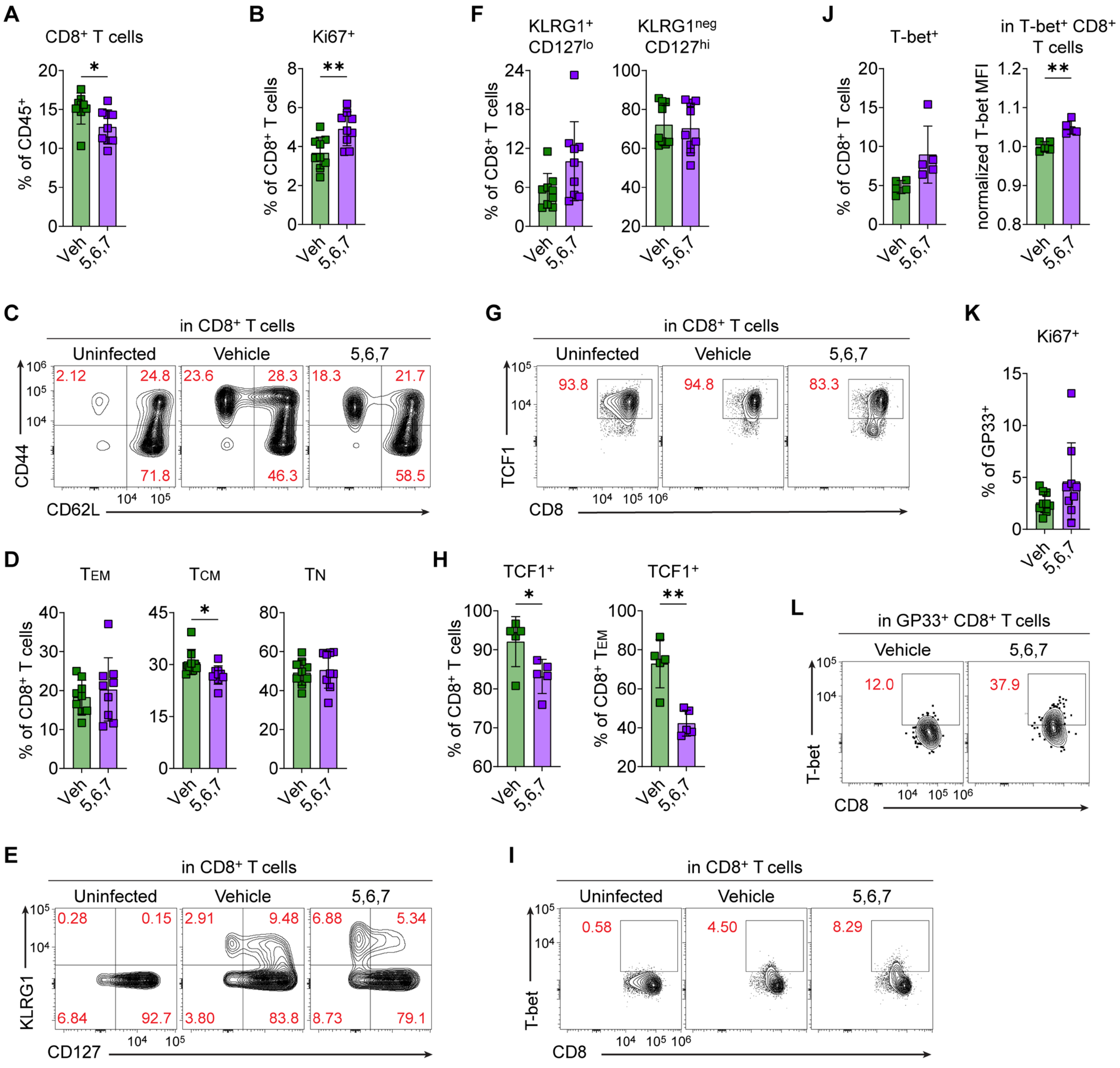


**Supplemental Figure S5. ATR inhibition during $CD8^+$ T cell clonal expansion permanently alters memory $CD8^+$ T cell phenotypes.**

Mice were infected with LCMV Armstrong on day 0 and were treated daily with Vehicle (Veh) or 75 mg/kg ATRi (days 5, 6, 7). Splenocytes were immune profiled by spectral flow cytometry at 7 weeks post infection. **A.** Quantitation of the $CD8^+$ T cells, as a percentage of total $CD45^+$ immune cells. **B.** Quantitation of the $Ki67^+$ $CD8^+$ T cells, as a percentage of total $CD8^+$ T cells. **C.** Representative contour plots showing CD44 and CD62L expression on $CD8^+$ T cells from uninfected control, Veh-treated, and ATRi-treated mice. **D.** Quantification of $CD8^+$ TEM ($CD44^+CD62L^{neg}$), TCM ($CD44^+CD62L^+$), and TN ($CD44^{neg}CD62L^+$), as percentages of total $CD8^+$ T cells. **E.** Representative contour plots showing KLRG1 and CD127 expression on $CD8^+$ T cells from uninfected control, Veh-treated, and ATRi-treated mice. **F.** Quantification of $KLRG1^+CD127^{lo}$ and $KLRG1^{neg}CD127^{hi}$ $CD8^+$ T cells, as percentages of total $CD8^+$ T cells. **G.** Representative contour plots showing TCF1 expression in $CD8^+$ T cells from uninfected control, Veh-treated, and ATRi-treated mice. **H.** Quantification of $TCF1^+$ $CD8^+$ T cells, as a percentage of total $CD8^+$ T cells, and quantitation of $TCF1^+$ $CD8^+$ TEM, as a percentage of total $CD8^+$ TEM. **I.** Representative contour plots showing T-bet expression in $CD8^+$ T cells from uninfected control, Veh-treated, and ATRi-treated mice. **J.** Quantification of T-$bet^+$ $CD8^+$ T cells, as a percentage of total $CD8^+$ T cells, and quantitation of T-bet median fluorescence intensity (MFI), normalized to the mean of Veh-treated mice, in T-$bet^+$ $CD8^+$ T cells. **K.** Quantitation of the $Ki67^+$ $GP33^+$ $CD8^+$ T cells, as a percentage of total $GP33^+$ $CD8^+$ T cells. **L.** Representative contour plots showing TCF1 expression in $GP33^+$ $CD8^+$ T cells from Veh-treated, and ATRi-treated mice. **A-F, K.** Data combined from two independent experiments with n = 4-5 mice per group for total n = 9 mice per group. Each experiment included one uninfected control. **G-J, L.** Data from one of the two above experiments, with n = 5 mice per group. **A-B, D, F, H, J-K.** Individual data points with mean ± SD bars shown. *p<0.05, **p<0.01, by unpaired, two-tailed Welch's t-test.

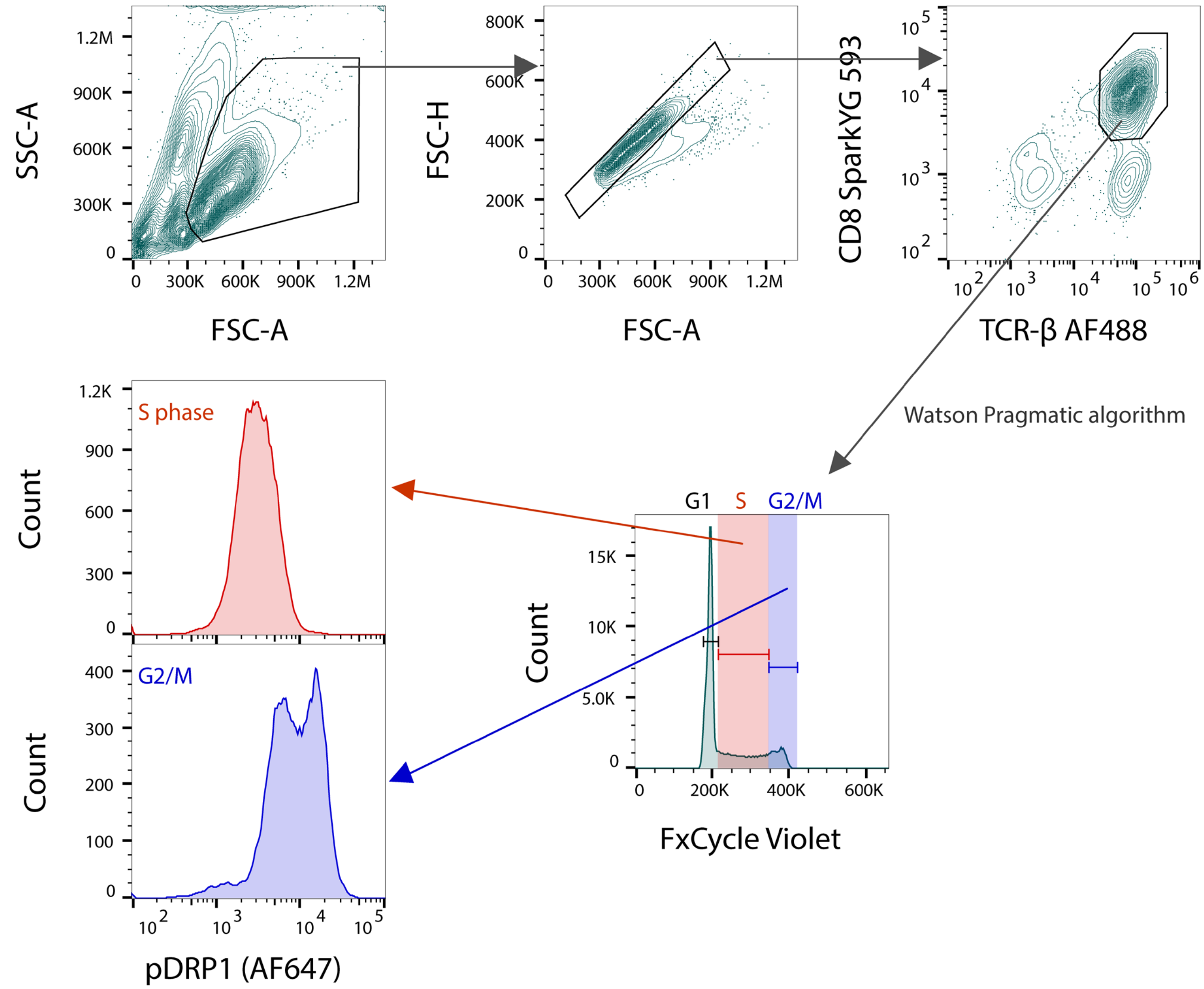


**Supplemental Figure S6. Gating strategy for analysis of pDRP1 in S and G2/M phase CD8⁺ T cells in pmel-1 splenocytes activated *ex vivo*.**

After any unstable portions of the run were excluded, cells of interest were gated based on FSC-A vs. SSC-A. Single cells were gated based on FSC-H vs. FSC-A and $CD8^+$ T cells ($CD8^+TCR\text{-}\beta^+$) were gated. Sub-G1 cells (if present) were excluded, and cell cycle phases were determined via univariate analysis of DNA content (FxCycle Violet) histograms using the Watson Pragmatic algorithm. Median fluorescence intensities (MFI) for pDRP1 were determined for cells in S phase and cells in G2/M phase.

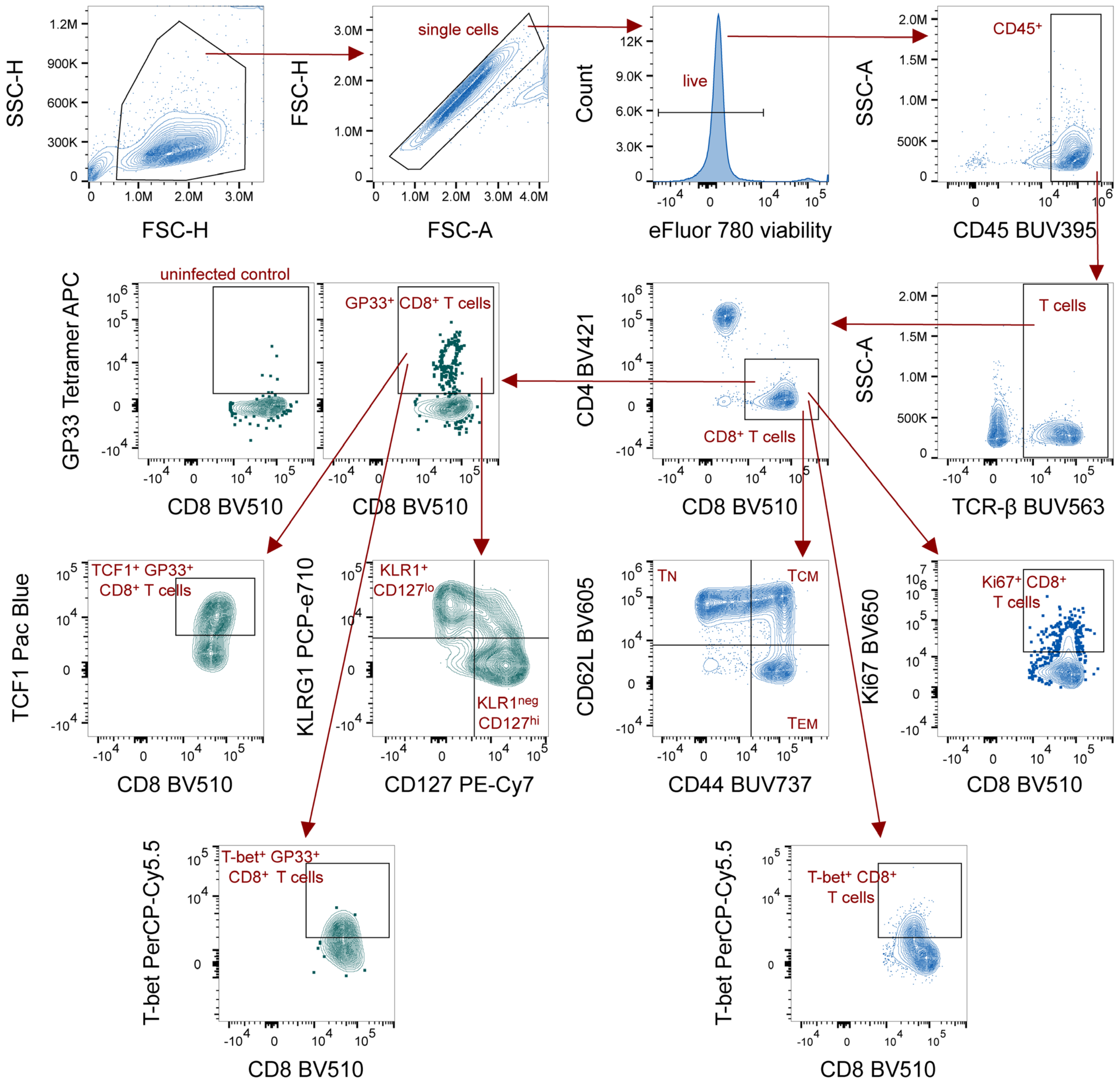


**Supplemental Figure S7. Gating strategy for immune profiling of LCMV-infected mice**

After any unstable portions of the run were excluded, cells of interest were gated based on FSC-H vs. SSC-H. Single cells were gated based on FSC-H vs. FSC-A, and live (viability dye negative) cells were selected. T cells were identified based on CD45 and TCR-β expression, CD8⁺ T cells were gated, and GP33-specific CD8⁺ T cells were identified based on GP33 Tetramer staining. Gating for GP33⁺ cells was determined using an uninfected control sample. Within both the CD8⁺ T cell pool and the GP33-specific $CD8^+$ T cell pool, cells were profiled for expression of Ki67, CD62L and CD44, KLRG1 and CD127, TCF1, and T-bet. Example plots are shown for each of these markers in either $CD8^+$ T cells or $GP33^+$ $CD8^+$ T cells. CD25, CD69, PD-1, and TIM-3 expression were also profiled but were not reported due to low signal, few positive events, or no differences between treatment groups. For day 18 immune profiling, when sufficient effector/effector memory $CD8^+$ T cells ($CD62L^{neg}CD44^+$, $T_{EM}$) were present, KLRG1 and CD127 expression were profiled within the $CD8^+$ $T_{EM}$ pool. Medium fluorescence intensities (MFI) for TCF1 and T-bet, within the $TCF1^+$ and T-$bet^+$ pools respectively, were quantitated.

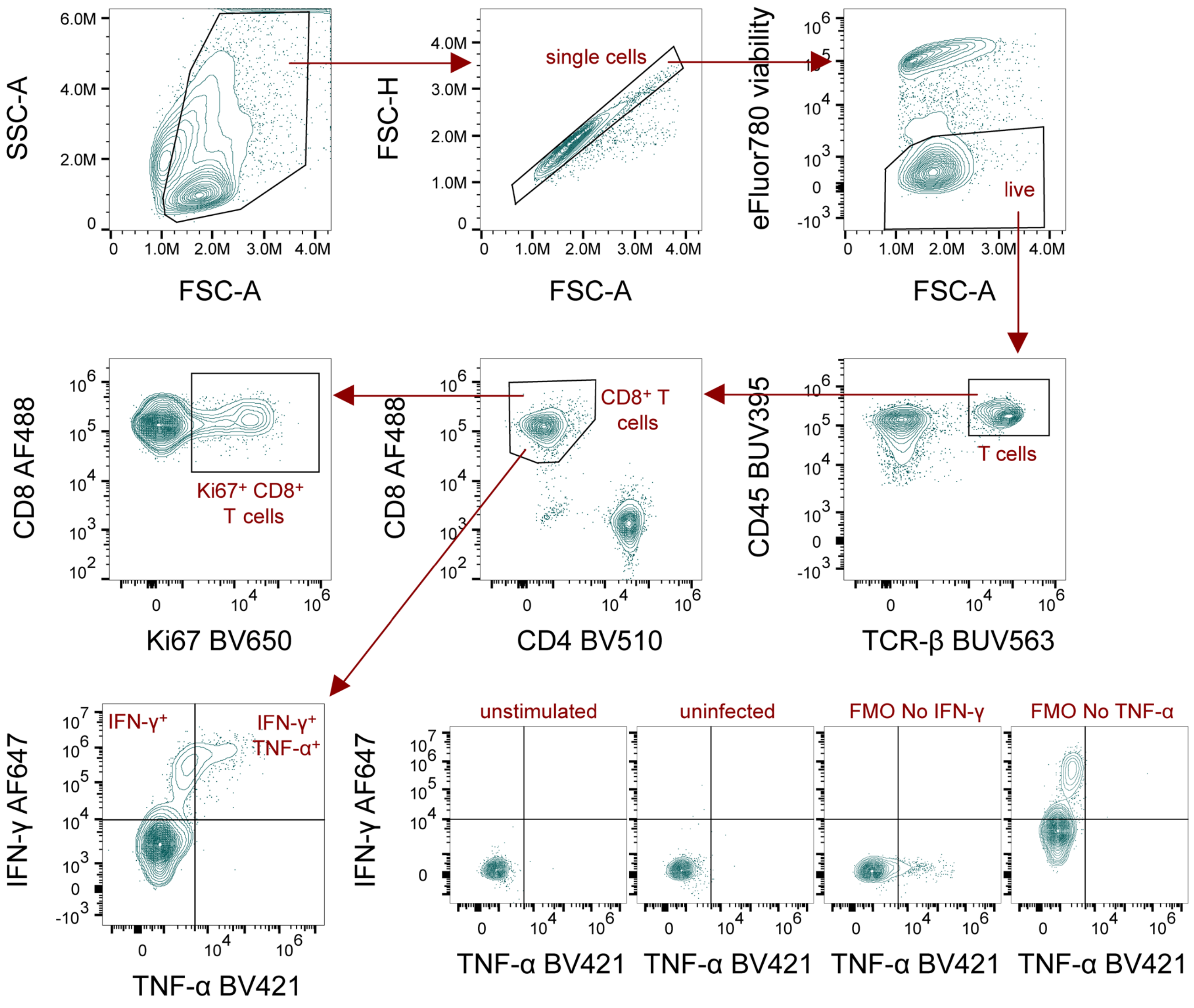


**Supplemental Figure S8. Gating strategy for cytokine and proliferation profiling in GP33-stimulated memory CD8$^{+}$ T cells.**

After any unstable portions of the run were excluded, cells of interest were gated based on FSC-A vs. SSC-A. Single cells were gated based on FSC-H vs. FSC-A, and live (viability dye negative) cells were selected. T cells were identified based on CD45 and TCR-β expression and CD8$^{+}$ T cells were gated. Within the CD8$^{+}$ T cell pool, cells were profiled for expression of Ki67, as well as IFN-γ and TNF-α. Example plots are shown for gating controls for IFN-γ and TNF-α, including unstimulated and uninfected control samples, as well as fluorescence minus one (FMO) controls.

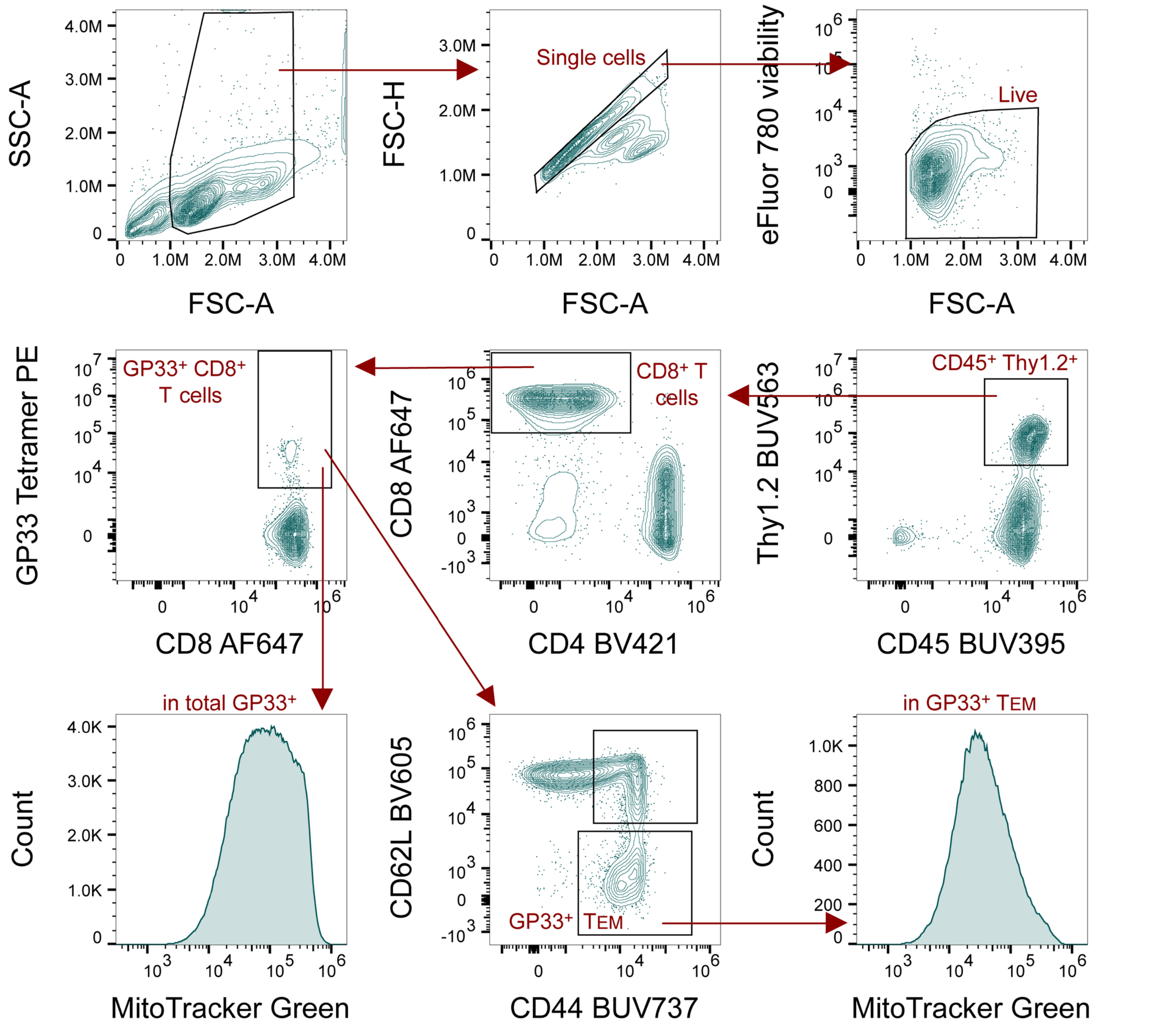


**Supplemental Figure S9. Gating strategy for analysis of mitochondrial mass in GP33-specific memory CD8⁺ T cells.**

After any unstable portions of the run were excluded, cells of interest were gated based on FSC-A vs. SSC-A. Single cells were gated based on FSC-H vs. FSC-A, and live (viability dye negative) cells were selected. T cells were identified based on CD45 and Thy1.2 expression, $CD8^+$ T cells were gated, and GP33-specific $CD8^+$ T cells were identified based on GP33 Tetramer staining. Gating for $GP33^+$ cells was determined using an uninfected control sample in addition to a fluorescence minus one (FMO) control (without GP33 Tetramer). GP33-specific $CD8^+$ T cells were then subset into $GP33^+$ central memory ($T_{CM}$, $CD62L^+CD44^+$) and $GP33^+$ effector memory ($T_{EM}$, $CD62L^{neg}CD44^+$) $CD8^+$ T cells. MitoTracker Green median fluorescence intensities (MFI) were quantified in the total $GP33^+$ $CD8^+$ T cell and the $GP33^+$ $CD8^+$ $T_{EM}$ populations.
